# A multi-step immune-competent genetic mouse model reveals phenotypic plasticity in uveal melanoma

**DOI:** 10.1101/2025.06.04.657841

**Authors:** Xiaonan Xu, Xiaoxian Liu, James J. Dollar, Xiao Liu, Neel Jasani, Benjamin Posorske, Sathya Neelature Sriramareddy, Vinesh Jarajapu, Jeffim N. Kuznetsoff, John Sinard, Richard L. Bennett, Jonathan D. Licht, Keiran S.M. Smalley, J. William Harbour, Xiaoqing Yu, Florian A. Karreth

## Abstract

Uveal melanoma (UM) is a highly aggressive intraocular malignancy with limited therapeutic options for metastatic disease. Existing transgenic UM mouse models inadequately recapitulate human disease progression, while transplant models lack immune competence for studying the tumor immune microenvironment and therapeutic interventions. To address these limitations, we developed a genetically engineered mouse model incorporating stepwise genetic alterations implicated in human UM progression. Spatiotemporally controlled expression of mutant GNAQ^Q209L^ from the endogenous locus induced choroidal nevi with limited penetrance. Concomitant BAP1 deletion enhanced nevus formation, while further MYC activation led to fully penetrant intraocular tumors with metastatic potential. Single-cell RNA sequencing revealed malignant cells segregated into Melanocytic and Neural Crest-like subpopulations characterized by distinct transcriptional and biosynthetic programs. Trajectory analyses inferred dedifferentiation from the Melanocytic toward the Neural Crest-like state during tumor progression. Comparison to human UM revealed commonalities with highly aggressive Class 2 UM, including gene expression signatures and copy number gains affecting genes that map to human chromosome 8q beyond the activated MYC allele, suggesting cooperative effects of multiple drivers in this chromosomal region. The tumor microenvironment featured immunosuppressive macrophage populations and exhausted T cells, closely resembling human UM. This physiologically relevant, immune-competent model provides a platform for investigating UM biology, functionally characterizing candidate driver genes, and developing immune-based therapeutic strategies.

**SIGNIFICANCE STATEMENT:** We developed a mouse model that resembles the genetic progression and phenotypic plasticity of human UM. This spatially controlled model confirms the critical role of driver mutations in GNAQ and BAP1, proposes MYC as a promoter of malignant transformation in coordination with other chromosome 8q genes, and reveals UM progression through distinct cellular states. This model offers an urgently needed preclinical platform for understanding the immunogenomics of UM and for testing immune and targeted treatments for this lethal cancer.

## INTRODUCTION

Uveal melanoma (UM) is a rare subtype of melanoma but the most common ocular malignancy (1). Approximately half of UM patients develop metastases, primarily in the liver, resulting in a median overall survival of less than one year (2,3). Currently, therapeutic options for metastatic UM patients remain highly limited (4), emphasizing the urgent need for a deeper understanding of the genetic and molecular mechanisms driving this disease to develop more effective treatments.

UM has a distinct mutational landscape compared to other melanoma subtypes, characterized by mutually exclusive activating hotspot mutations in *GNAQ* or *GNA11*, which encode Gα subunits of heterotrimeric G-proteins (5). Notably, *GNAQ/11* mutations are also found in choroidal nevi, yet only about 1 in 8,800 of these nevi progress to UM (6,7), indicating that additional genetic alterations are necessary for malignant transformation. Most UM undergo subsequent nearly mutually exclusive prognostic mutations in *BAP1*, *SF3B1* or *EIF1AX*, associated with high, intermediate and low metastatic risk, respectively. Additionally, canonical chromosome copy number alterations (CNA) are common, including monosomy 3, gains of 6p and 8q, and losses of 1p, 6q and 8p (8). While very little is understood about the role of these chromosomal aberrations in UM, it has been speculated that the oncogene MYC, located at chromosome 8q24, may play a role in UM progression (9).

UM can be classified into distinct prognostic subtypes based on a 15-gene expression profile (15-GEP) with Class 1 tumors being associated with *EIF1AX* and *SF3B1* mutations and low to intermediate metastatic rate, while Class 2 tumors are associated with *BAP1* loss-of-function mutations and a high propensity for metastasis (10). Chromosome 8q gains occur in both Class 1 and Class 2 UM, but the number of extra copies of 8q tends to be greater in Class 2 tumors, correlating with increased metastatic potential (11).

This landscape of genetic alterations provides a foundation for developing genetically engineered mouse models (GEMMs) that can facilitate preclinical UM research and to serve as platforms for testing therapeutic strategies. Inducible alleles of mutant *GNAQ* (12) and *GNA11* (13) have been developed, and when crossed to melanocyte-specific Cre mice, they induce rapid ocular melanoma formation (12,13). These findings establish *GNAQ/11* as initiating oncogenes in UM. However, single mutant *GNAQ/11* mice develop UM within weeks, in contrast to the slower progression seen in human patients, where cooperating alterations are required. This discrepancy reflects that these models do not fully recapitulate the genetics and biology of human UM, limiting their utility for studying UM progression. Possible reasons for this discrepancy include inappropriate timing of mutant *GNAQ/11* activation during embryonic development and non-physiological expression levels from ectopic promoters. Moreover, these models develop undesired cutaneous melanomas, precluding long-term studies aimed at modeling metastatic progression following enucleation of the primary tumor. These limitations underscore the need for more refined GEMMs that better reflect human UM development.

In this study, we report the development of an inducible *Gnaq*-mutant allele expressed at physiological levels from its endogenous promoter. Restricting mutant *Gnaq* activation to the eye resulted in choroidal nevi, and concomitant *BAP1* deletion enhanced the penetrance of nevus formation. The addition of an activated *MYC* allele, a candidate oncogene located on chromosome 8q24, led to fully penetrant intraocular tumor formation with features of human UM. Cell lines derived from this triple-mutant model formed tumors in orthotopic transplants and exhibited metastasis. Furthermore, scRNAseq of murine tumors revealed intra-tumoral phenotypic heterogeneity, as well as inferred copy number alterations that correspond to many of the genes lost on chromosome 3 and gained on chromosome 8q in human UM. Cross-species comparison confirmed similarities to human Class 2 UM, reinforcing the relevance of this model for studying disease progression. This UM model will facilitate preclinical research into the pathogenesis of UM and enable the identification of potential drivers and vulnerabilities.

## RESULTS

### Generation of an inducible, physiologically expressed Gnaq^Q209L^ allele

To generate a mouse model that faithfully recapitulates the genetics and biology of human UM, we used a “mini-gene” approach. To this end, we replaced exon 5 of the *Gnaq* gene with a loxP-flanked cDNA cassette containing wildtype exons 5-7 followed by exon 5 mutated to encode the hotspot Q209L mutation. This allele, termed Gnaq^CA^, expresses wildtype GNAQ until Cre-mediated excision of the cDNA cassette enables splicing into mutant exon 5, resulting in expression of GNAQ^Q209L^ from its endogenous locus (**Fig. 1A**). To validate functionality and physiological expression levels of the Gnaq^CA^ allele, we established mouse embryonic fibroblasts (MEFs) from Gnaq^CA/+^ mice followed by infection with adenoviral Cre. Sanger sequencing of Gnaq exon 5 from Cre-infected Gnaq^CA/+^ MEFs confirmed heterozygous expression of the mutant allele (**Fig. 1B**). Importantly, Cre administration led to the hyperactivation of Gq-dependent pathways, as shown by increased levels of activating ERK phosphorylation and decreased levels of inactivating YAP phosphorylation, in the absence of changes to the total expression levels of GNAQ (**Fig. 1C**). Thus, the Gnaq^CA^ allele enables inducible expression of mutant GNAQ^Q209L^ at physiological levels, aiming to model the GNAQ activation observed in human UM patients.

**Figure 1.**
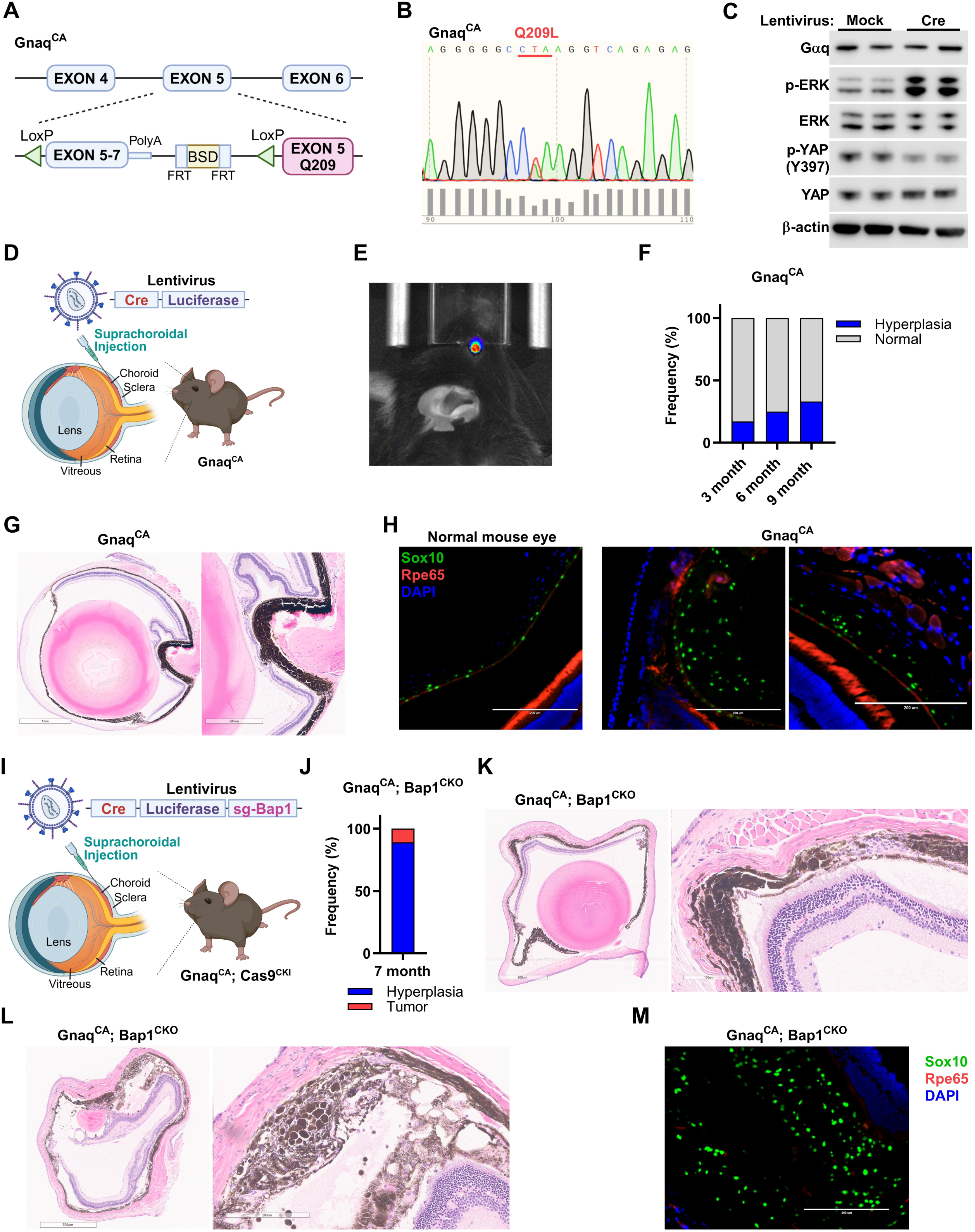
Generation of a genetically engineered mouse model of choroidal nevi driven by *Gnaq* mutation and *Bap1* loss. **A,** Schematic diagram of the Gnaq conditional activation (Gnaq^CA^) allele. **B,** Mouse embryonic fibroblasts (MEFs) isolated from Gnaq^CA^ mice were infected by lentiviral Cre. PCR amplification and Sanger sequencing of Gnaq exon 5 from cDNA confirmed expression of the Q209L mutation. **C,** Western blot of Cre-infected Gnaq^CA^ MEFs revealed changes in Gαq downstream signaling but physiological GNAQ expression. **D,** Schematic diagram of Gnaq conditional activation (Gnaq^CA^) in the uvea by in situ lentiviral delivery of Cre and Luciferase (pL-CL). **E,** Representative in vivo bioluminescence imaging of a Gnaq^CA^ mouse infected with pL-CL. **F,** Quantification of frequency of choroidal hyperplasia in Gnaq^CA^ mice at the indicated timepoints. **G,** Representative image of H&E staining showing localized choroidal hyperplasia in a Gnaq^CA^ mouse. **H,** Representative fluorescent IHC showing SOX10 and RPE65 expression in uveal tracts of wildtype and Gnaq^CA^ mice. **I,** Schematic diagram of Gnaq conditional activation and Bap1 conditional CRISPR knockout (Gnaq^CA^; Bap1^CKO^) in the uvea by in situ lentiviral delivery of Cre, Luciferase, and sgBap1 (pL-CLB). **J,** Quantification of choroidal hyperplasia and tumors in Gnaq^CA^; Bap1^CKO^ mice at 7 months. **K** and **L,** Representative images of H&E staining of choroidal hyperplasia (**K**) and micro-tumor (**L**) in Gnaq^CA^; Bap1^CKO^ mice. **M**, Representative fluorescent IHC showing SOX10 and RPE65 expression in the uveal tract of a Gnaq^CA^; Bap1^CKO^ mouse.

### Melanocyte-specific activation of GNAQ^Q209L^

To investigate the consequences of GNAQ^Q209L^ activation in melanocytes, we crossed Gnaq^CA^ mice to a mouse strain that expresses tamoxifen-inducible Cre from the melanocyte-specific Tyrosinase promoter (Tyr-CreERt2)(14). All compound mutant Gnaq^CA^; Tyr-CreERt2 mice developed extensive skin hyperpigmentation, especially on non-hair bearing skin, as well as multiple nevi in the absence of Tamoxifen administration at 2-3 months of age (**Supplementary Fig. S1A**). Several nevi per mouse progressed to rapidly growing pigmented or unpigmented skin tumors at 4-6 months of age (**Supplementary Fig. S1B,C**). Leakiness of the Tyr-CreERt2 allele had been noted previously in the Braf^V600E^; Pten^FL/FL^ background; however, the extent and kinetics of tumorigenesis in Gnaq^CA^; Tyr-CreERt2 mice and the fact that melanomas developed in the absence of cooperating mutations was unexpected. The skin phenotype in Gnaq^CA^; Tyr-CreERt2 mice occurred before overt ocular abnormalities were observed. When the skin disease required euthanasia, we assessed ocular histology and identified extensive pigmented hyperplasia of the entire uveal layer (**Supplementary Fig. S1D**). SOX10 (marker for melanocytes and retinal pigment epithelium) and RPE65 (marker specific for retinal pigment epithelium) staining revealed that uveal hyperplasia was composed of melanocytes, not retinal pigmented epithelium cells (**Supplementary Fig. S1E**). These findings demonstrated that activation of the Gnaq^CA^ allele in uveal melanocytes robustly induces hyperplasia. However, similar to previous mutant Gnaq or Gna11 strains (12,13), the extensive skin disease precludes the use of melanocyte-specific Cre strains to study GNAQ^Q209L^-driven uveal melanoma.

### Development of a uveal melanoma mouse model

To restrict expression of GNAQ^Q209L^ to the eye and circumvent recombination of the Gnaq^CA^ allele in cutaneous or organ-resident melanocytes, we generated a lentivirus containing Cre and Luciferase (pL-CL) (**Fig. 1D**). The virus was directly delivered to the uveal tract via suprachoroidal injection (**Fig. 1D**) and successful in situ infection was confirmed by detecting Luciferase activity through in vivo bioluminescence imaging (**Fig. 1E**). Histological evaluation of eyes from Gnaq^CA^ mice at 3, 6, and 9 months post-infection revealed localized choroidal thickening with increasing penetrance, occurring in 33% of mice at the 9-months timepoint (**Fig. 1F,G**). Hyperplasia of choroidal melanocytes was evident, without violation of Bruch’s membrane or any effect on the overlying retinal pigment epithelium. Importantly, this approach completely prevented any undesired formation of cutaneous lesions and tumors. SOX10 and RPE65 staining showed an increased number of melanocytes in choroidal hyperplasia (**Fig. 1H**), validating that lentiviral delivery of Cre to activate GNAQ^Q209L^ expression results in hyperplasia of choroidal melanocytes. Thus, this approach enables spatiotemporal control over GNAQ^Q209L^ activation and results in a phenotype that resembles choroidal nevus formation in humans.

To generate a versatile platform in which secondary genetic hits can be readily introduced, we bred Gnaq^CA^ mice to R26-LSL-Cas9 (15) mice. Double mutant mice were suprachoroidally infected with the pL-CL virus to which a sgRNA targeting *Bap1* was added (pL-CLB). *BAP1* is considered a key tumor suppressor in human UM, and this approach enables CRISPR-mediated knockout of *Bap1* in situ in the context of GNAQ^Q209L^ activation (Gnaq^CA^; Bap1^CKO^) (**Fig. 1I, Supplementary Fig. S1F,G**). Notably, all 9 Gnaq^CA^; Bap1^CKO^ mice exhibited localized hyperplasia 7 months after lentiviral administration (**Fig. 1J,K**). Interestingly, while *Bap1* deletion increased the penetrance of choroidal hyperplasia, it did not efficiently promote UM formation. Indeed, we observed only a single Gnaq^CA^; Bap1^CKO^ mouse bearing a small choroidal tumor (**Fig. 1L**). Cells in this small tumor were SOX10 positive and RPE65 negative, consistent with a choroidal melanoma (**Fig. 1M**). Thus, concurrent GNAQ^Q209L^ activation and Bap1 deletion promotes fully penetrant choroidal nevus formation with rare progression to UM, suggesting that additional genetic events may be required in this model system to drive UM development with high penetrance.

Progressive gain of copies of chromosome 8q occur throughout the evolution of UM, starting in low grade Class 1 tumors and increasing in copy number in Class 2 UM and metastatic tumors (16). *MYC* is located on chromosome 8q, overexpressed upon 8q gain (17), and thus one candidate for the oncogene(s) targeted for upregulation by8q gain. Moreover, during UM progression. We therefore selected MYC as a candidate oncogene on 8q and tested whether its expression can cooperate with *GNAQ* mutation and *BAP1* loss to enhance malignant transformation and promote UM formation. To test this, we crossed Gnaq^CA^; R26-LSL-Cas9 mice to a strain harboring a Cre-dependent stabilized *MYC* allele (Myc^T58A^) (18). Suprachoroidal infection of compound mutant mice with the pL-CLB lentivirus generated Gnaq^CA^; Bap1^CKO^; Myc^CKI^ mice having three UM-associated genetic alterations in their uveal melanocytes (**Fig. 2A**). Notably, Gnaq^CA^; Bap1^CKO^; Myc^CKI^ mice developed aggressive UM with short latency and full penetrance, resulting in a median survival of ∼3 months (**Fig. 2B**). In vivo bioluminescence imaging identified strong Luciferase signal from primary tumors (**Fig. 2C**), enabling monitoring of primary tumor growth and, potentially, in vivo tracking of metastasis formation. At endpoint, intraocular tumors filled the vitreous cavity (**Fig. 2D**), leading to significant proptosis (**Supplementary Fig. S2A**). We also observed extraocular tumor extension in several eyes, consistent with locally invasive disease (**Supplementary Fig. S2A**). Intraocular tumors grew into the sub-retinal space, displacing the retina and filling the posterior chamber. Histologically, the tumors were largely unpigmented, although rare foci of pigmentation were frequently observed. Some cases exhibited zones of necrosis with corresponding variable inflammatory responses. The tumors grew as sheets of poorly differentiated epithelioid cells with occasional nesting. Cytologically, tumors cells varied in size, with round to oval nuclei containing one or two prominent nucleoli. The cytoplasm appeared somewhat granular, and some tumors displayed areas of partial cytoplasmic clearing (**Fig. 2E**). IHC analysis demonstrated RPE65 negativity and clonal SOX10 positivity (**Fig. 2F**), while Periodic Acid-Schiff staining revealed a PAS negative pattern (**Fig. 2G**). The histologic and marker analyses suggest that the intraocular tumors in Gnaq^CA^; Bap1^CKO^; Myc^CKI^ mice may represent a mouse equivalent of human UM.

**Figure 2.**
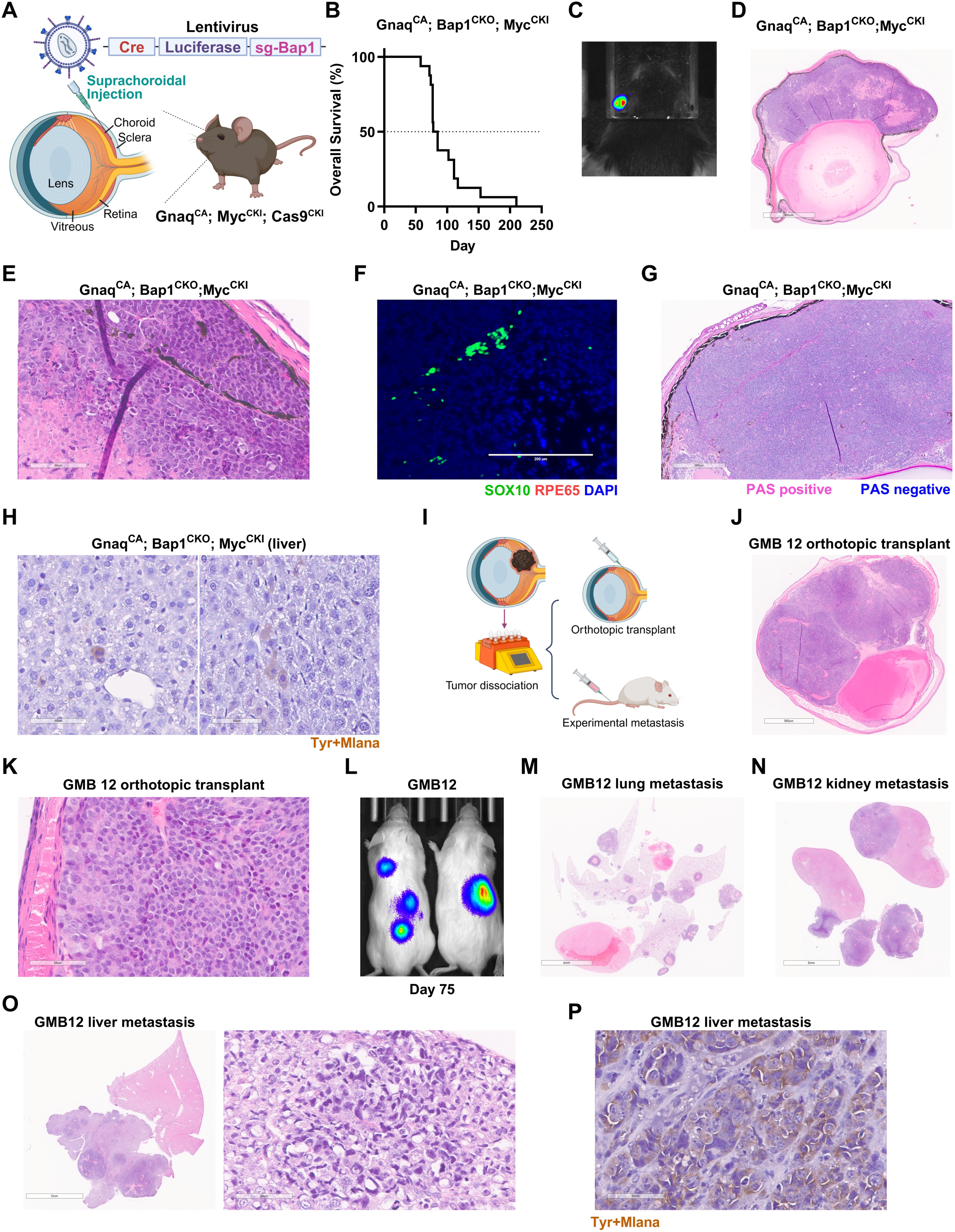
MYC promotes UM formation. **A,** Schematic diagram of Gnaq conditional activation, Bap1 conditional CRISPR knockout, and Myc conditional knock-in (Gnaq^CA^; Bap1^CKO^; Myc^CKI^) in the uvea by in situ delivery of pL-CLB. **B,** Kaplan-Meier overall survival curve of Gnaq^CA^; Bap1^CKO^; Myc^CKI^ mice after pL-CLB administration. **C,** Representative in vivo bioluminescence imaging showing strong signal from a primary intraocular tumor. **D** and **E,** Representative images of H&E staining of intraocular tumors from Gnaq^CA^; Bap1^CKO^; Myc^CKI^ mice. **F,** Representative fluorescent IHC showing SOX10 and RPE65 expression in a tumor from a Gnaq^CA^; Bap1^CKO^; Myc^CKI^ mouse. Tumors are clonally positive for SOX10 and negative for RPE65 **G,** Representative image of Periodic acid–Schiff (PAS) staining of a tumor from a Gnaq^CA^; Bap1^CKO^; Myc^CKI^ mouse. Tumors displayed minimal PAS positivity **H,** Representative images of IHC using a melanoma cocktail (Tyrosinase + Melan-A) to detect disseminated UM cells in the livers of Gnaq^CA^; Bap1^CKO^; Myc^CKI^ mice. **I,** Schematic diagram of orthotopic and metastatic transplant of Gnaq^CA^; Bap1^CKO^; Myc^CKI^ primary tumor cells in NSG mice. **J** and **K,** Representative images of H&E staining of orthotopic transplant of Gnaq^CA^; Bap1^CKO^; Myc^CKI^ primary tumor cells (GMB12). **L,** Representative in vivo bioluminescence imaging showing signal from GMB12 cells 75 days after tail vein inoculation. **M-O,** Representative images of H&E staining of metastasis in lung (**M**), kidney (**N**), and liver (**O**). **P,** Representative IHC using a melanoma cocktail (Tyrosinase + Melan-A) on the liver of a mouse injected with GMB12 cells.

### Metastatic uveal melanoma models

Because *BAP1* loss-of-function and 8q gains are associated with increased risk of UM metastasis, we analyzed livers of Gnaq^CA^; Bap1^CKO^; Myc^CKI^ mice. We observed no overt metastasis in any of the analyzed livers. However, IHC for a melanocyte marker cocktail (TYR + MLANA) identified rare positive cells in livers from two out of seven Gnaq^CA^; Bap1^CKO^; Myc^CKI^ mice, which could represent disseminated tumor cells (**Fig. 2H, Supplementary Fig. S2B**). Because the rapid growth of the primary intraocular tumors precludes long term metastasis analysis, we tested the metastatic potential of tumor cells derived from Gnaq^CA^; Bap1^CKO^; Myc^CKI^ mice in transplant models (**Fig. 2I, Supplementary Fig. S2C**). Orthotopic transplant tumors inoculated via suprachoroidal injection of mouse tumor cells exhibited a similar histology to primary tumors (**Fig. 2J,K**) and maintained a RPE65 negative and clonal SOX10 positive pattern (**Supplementary Fig. S2D**). Furthermore, tail vein inoculation of mouse tumor cells resulted in metastases to the lungs, kidney, and liver (**Fig. 2L-O**). Metastases maintained the histology of the primary tumor (**Fig. 2M-O**, **Supplementary Fig. S2E,F**) and were positive for melanoma markers (**Fig. 2P**, **Supplementary Fig. S2G**). We then derived cell lines from liver and lung metastasis after repeated in vivo passage to select for tumor cells with liver and lung organotropism (**Supplementary Fig. S2H,I**). When cultured in vitro, liver-tropic cells adopted an amoeboid morphology while lung-tropic cells exhibited a more mesenchymal appearance (**Supplementary Fig. S2H,I**). Thus, expression of GNAQ^Q209L^, loss-of-function of BAP1, and activation of MYC promotes the formation of intraocular tumors resembling human UM with metastatic potential.

### Phenotypic characterization of murine uveal melanoma by single cell RNA sequencing

We subjected three tumors from Gnaq^CA^; Bap1^CKO^; Myc^CKI^ mice to single cell RNA sequencing (scRNA-seq) to characterize the cellular composition and phenotypic landscape of murine uveal melanomas (**Supplementary S3A**). After quality control and doublets removal (**Supplementary Fig. S3B**), 24,455 cells were taken forward for downstream analysis. Approximately 75% of cells in each tumor were classified as malignant (**Fig. 3A, Supplementary S3A-D**). In addition to malignant cells, all three tumors exhibited immune cell infiltration, with the greatest contribution to the immune cell pool coming from macrophages/monocytes and neutrophils, with the former classified as *Arg*1+ macrophages (*Arg1*, *Ccl9*, *Flrt3*) and *C1qa*+ macrophages (*C1qa*, *C1qb*, *C1qc*, *Fcrls*) (**Fig. 3A, Supplementary S3C,D**). T/NK cells and B cells were less abundant in all three tumors (**Fig. 3A, Supplementary S3C**). Examining immune cell populations revealed the expression of several immune exhaustion markers, including *Lag3* expression in T cells and *Harvk2* expression in mature regulatory dendritic cells (mregDC), monocytes, and macrophages, consistent with previous reports of T cell states within the Class 2 UM immune microenvironment (19) (**Supplementary Fig. S3E**). Cancer-associated fibroblasts (CAF) were also present in all three tumors, and inflammatory CAF (*Cxcl5*, *Il11*, *Saa3*), Myofibroblast-like CAF (*Actg2*, *Cd248*, *Myh11*), and activated CAF were further identified (**Fig. 3A, Supplementary Fig. S3C-D**). Other normal stromal cells including retinal cells (photoreceptor cells, bipolar cells, Müller glial cells), retinal pigmented epithelium (RPE) cells, multiciliated epithelial cells, and endothelial cells were also detected (**Fig. 3A, Supplementary Fig. S3C-D**).

**Figure 3.**
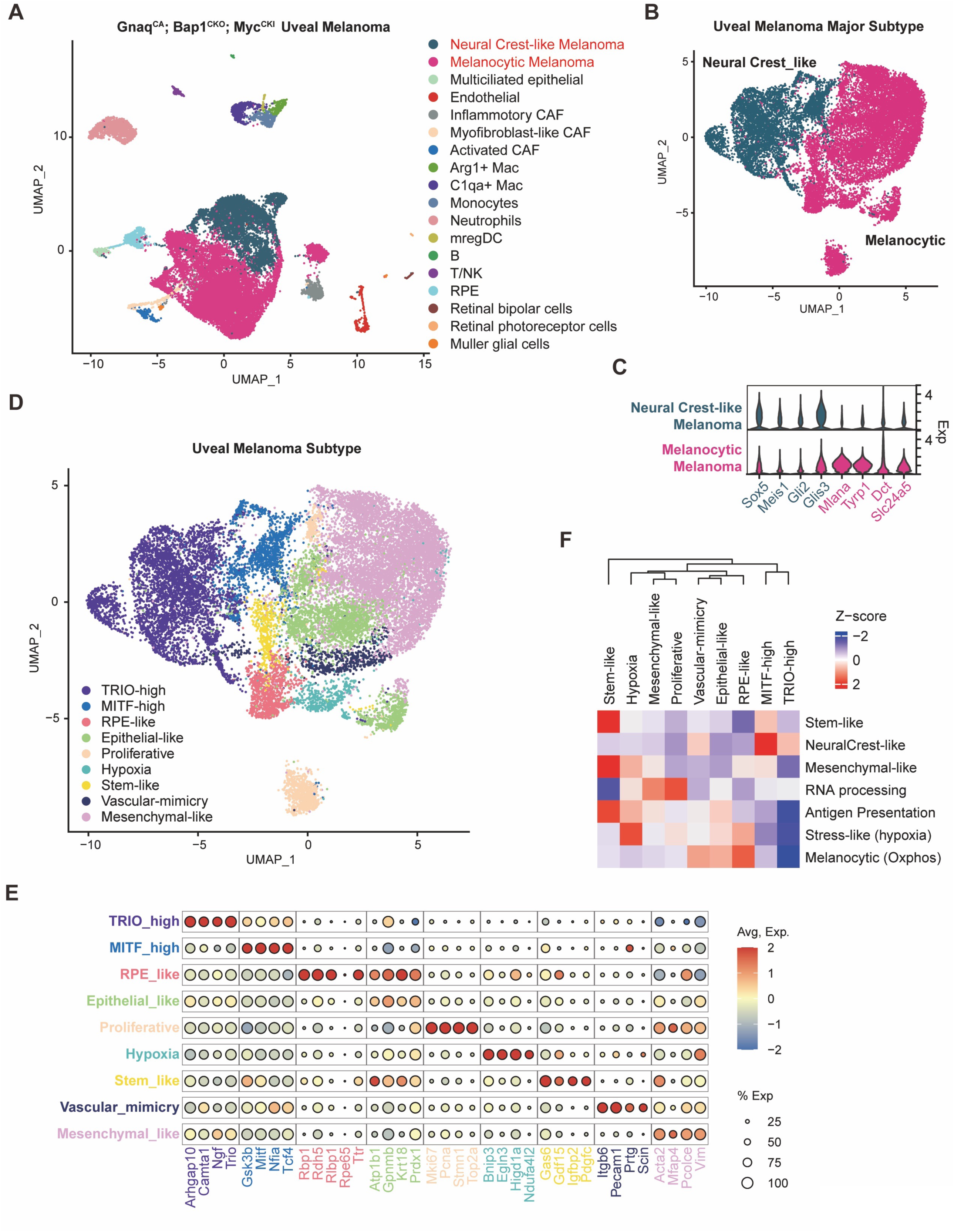
Characterization of phenotypic diversity in murine UM. **A,** UMAP plot showing malignant and microenvironmental cell types in UMs from Gnaq^CA^; Bap1^CKO^; Myc^CKI^ mice analyzed by scRNA-seq. Cell types are color-coded. **B,** UMAP plot showing the two major subtypes in UMs from Gnaq^CA^; Bap1^CKO^; Myc^CKI^ mice, Melanocytic and Neural Crest-like (color-coded). **C,** Violin plot showing the expression of Melanocytic and Neural Crest-like subtype markers. **D,** UMAP plot showing phenotypic clusters among the malignant UM cells from Gq^CA^; Myc^CKI^; Bap1^CKO^ tumors. Clusters are color-coded. **E,** Dot plot showing the expression of cluster markers, with dot color indicating the average expression levels and dot size indicating the percentage of cells in each cluster expressing the respective marker. **F,** Heatmap showing the relationship of murine UM cell states to murine cutaneous melanoma phenotypic states based on the relative expression of the indicated gene signatures.

The malignant cell population could be broadly categorized into two major subtypes in the UMAP space, Melanocytic and Neural Crest-like (**Fig. 3B**). The Melanocytic subtype was defined by high expression of *Tyrp1*, *Dct*, *Mlana*, and *Slc24a5*, representing a developmental stage of mature melanocytes (**Fig. 3C, Supplementary Fig. 3D**). The Neural Crest-like subtype was defined by expression of *Gli2*, *Glis3*, *Meis1*, and *Sox5*, representing developmental stages of neural crest progenitors, premelanoblasts, and melanoblasts. Analysis of marker gene expression enabled further sub-categorization of cell populations in the Melanocytic and Neural Crest-like major subtypes into minor subtypes of tumor cells (**Fig. 3D,E**). This led to the identification of TRIO-high (*Trio, Ngf, Arhgap10, Camta1*) and MITF-high (*Mitf, Gsk3b, Nfia, Tcf4*) clusters in the Neural Crest-like major subtype, which represent neural crest cells and early melanoblasts at developmental stages, respectively. Moreover, Epithelial-like (*Krt18, Atp1b1, Gpnmb, Prdx1*), Mesenchymal-like (*Vim, Acta2, Mfap4, Pcolce*), Proliferative (*Mik67, Pcna, Stmn1, Top2a*), Hypoxia (*Bnip3, Egln3, Higd1a, Ndufa4l2*), Stem-like (*Gas6, Gdf15, Igfbp2, Pdgfc*), and RPE-like (*Rbp1, Rdh5, Rlbp1, Ttr*) clusters were identified in the Melanocytic major subtype. A vascular-mimicry cluster (*Itgb6, Pecam1, Prtg, Scin*), a cell state that has been associated with Class 2 GEP (20), chromosome 3 loss and 8q gain (21), contains both Neural Crest-like and Melanocytic cells (**Fig. 3D,E**). Both the Neural Crest-like and Melanocytic major subtypes as well as the nine minor subtypes were equally represented in all three intraocular tumors (**Supplementary Fig. S4A,B**) and show similar quality control metrics (**Supplementary Fig. S4C**). The phenotypic categorization of murine tumor cells partially matched the phenotypic states observed in murine cutaneous melanoma on a Nras^Q61K^; p16Ink4a^-/-^ background (22), indicating that despite being induced by distinct driver mutations, mouse models of uveal and cutaneous melanoma may share some phenotypic and molecular commonalities (**Fig. 3F, Supplementary Fig. S4D**). These analyses revealed a complex tumor microenvironment with malignant cells exhibiting various phenotypic states and immune microenvironment reminiscent of that seen in human UM.

### Melanocytic and Neural Crest-like UM cells exhibit distinct gene expression profiles

Given our finding that MYC promotes intraocular tumor development in this murine model of UM, we further examined the expression of *Myc* and its target genes. The expression of endogenous *Myc* was moderately higher in Melanocytic cells compared to Neural Crest-like cells (**Fig. 4A,B**), and the expression of ectopic *Myc* (using Luciferase, which is linked to Myc^T58A^ in the construct, as a surrogate for transgene expression) was lost in a subset of Neural Crest-like cells (**Supplementary Fig. S5A**). Endogenous *Myc* expression was not significantly changed among the two and seven minor subtypes that constitute the Neural Crest-like and Melanocytic major subtypes, respectively (**Supplementary Fig. S5B**). These findings indicate that the Neural Crest-like and Melanocytic states differentially rely on MYC activity. Accordingly, the expression of MYC targets (defined by the GSEA_Hallmark geneset) is significantly higher in Melanocytic than Neural Crest-like cells (**Fig. 4C**). Gene set enrichment analysis of Hallmark signatures identified enrichment of MYC targets, as well as oxidative phosphorylation, reactive oxygen species (ROS) pathway, mTORC1 signaling, UV response, fatty acid metabolism, and apoptosis in Melanocytic UM cells (**Fig. 4D**). Gene ontology (GO) analysis showed that ribosome and mitochondrial signatures are enriched in Melanocytic UM cells (**Fig. 4E**). Notably, among the genes exhibiting the highest positive correlation with *Myc* expression were a number of genes encoding ribosomal proteins, and this expression correlation is conserved in human UM (**Fig. 4F**). Indeed, the expression of 84 of 90 ribosome genes and 74 of 78 mitochondrial ribosome genes is increased in the Melanocytic state compared to Neural Crest-like state (**Fig. 4G**). These findings indicate that Melanocytic and Neural Crest-like UM subtypes display different molecular features that correlate with *Myc* expression.

**Figure 4.**
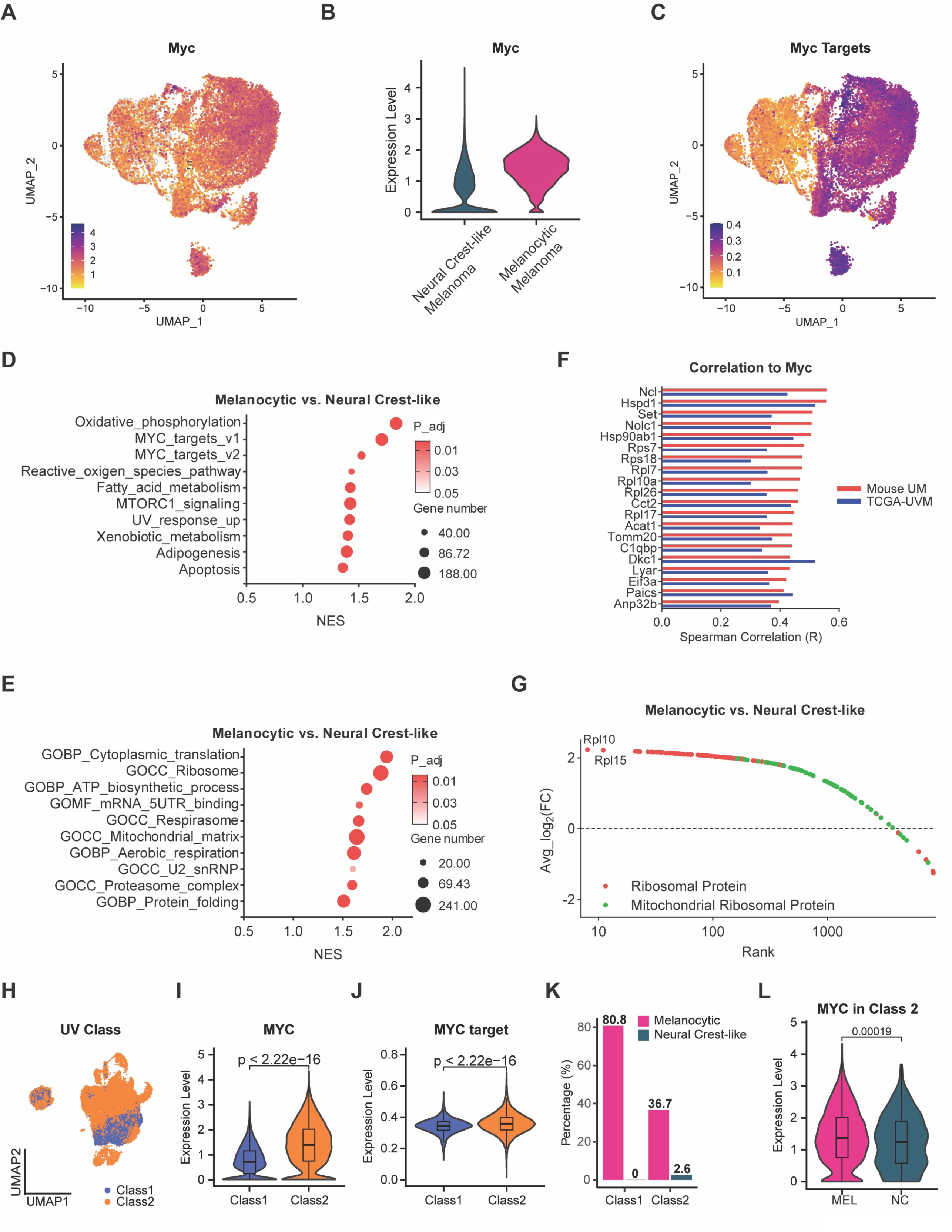
MYC activity across the UM progression continuum. **A,** UMAP plot showing *Myc* expression in malignant UM cells from Gnaq^CA^; Bap1^CKO^; Myc^CKI^ tumors. **B,** Violin plots comparing endogenous *Myc* expression between the Melanocytic and Neural Crest-like subtypes. **C,** UMAP plot showing expression of Myc targets in murine UM cells, color coded by average expression of canonical Myc targets from “GSEA_Hallmark_MYC_targets_v1/v2”. **D** and **E,** Gene set enrichment analysis (GSEA) of Hallmark signature (**D**) and gene ontology (GO) (**E**) in Melanocytic and Neural Crest-like UM cells. **F,** Spearman correlation between *Myc* expression and the indicated genes in murine UM from the scRNA-seq analysis and human UM from the UM dataset form TCGA (TCGA-UVM). **G,** Scatter plot showing the expression changes of 90 ribosome genes and 78 mitochondrial ribosome genes between Melanocytic and Neural Crest-like cells. **H,** UMAP of human UM scRNA-seq dataset, color coded by Class 1 and Class 2 primary tumors. **I** and **J,** Violin plots showing expression of *MYC* (**I**) and *MYC* targets (**J**) in Class 1 and Class 2 human UM. **K,** Percentage of Class 1 and Class 2 human UM cells positive for the Melanocytic or Neural Crest-like signatures. **L,** Violin plot showing *MYC* expression in Class 2 human UM cells positive for the Melanocytic or Neural Crest-like signatures.

We then determined whether the molecular features identified in our murine model of UM are conserved in human Class 1 or Class 2 primary UM by analyzing a previously published scRNA-seq dataset (19) (**Fig. 4H**). Notably, *MYC* is expressed at significantly higher levels in Class 2 UM cells than Class 1 UM cells (**Fig. 4I, Supplementary Fig. S5C**), with a concurrent increase in MYC target gene expression (**Fig. 4J, Supplementary Fig. S5D**). Moreover, human UM expressed gene signatures defining the murine Melanocytic and Neural Crest-like states; however, there were stark differences between Class 1 and Class 2 UMs. Class 1 UMs predominantly express the Melanocytic signature but are negative for the Neural Crest-like signature (**Fig. 4K, Supplementary Fig. S5E**). Conversely, a smaller proportion of Class 2 UM cells are positive for the Melanocytic signature, and cells in the Neural Crest-like state appeared (**Fig. 4K, Supplementary Fig. S5E**). Both Class 1 and Class 2 UMs contain cells that are positive for both signatures (**Fig. 4K, Supplementary Fig. S5E**), but whether this represents cells transitioning from one state to the other remains to be determined. Similar to the findings in our mouse model, Class 2 UM cells in the Neural Crest-like state express significantly lower *MYC* levels than Melanocytic cells (**Fig. 4L**).

Four of the genes in the 15-GEP prognostic test – *CDH1, ECM1, HTR2B,* and *RAB31* – are differentially overexpressed in Class 2 UM (24258991, 39052972), and three of these genes – *Htr2b, Ecm1,* and *Rab31* – are significantly upregulated in murine Neural Crest-like UM cells, consistent with the expression profile of *BAP1*-deficient Class 2 UM (**Supplementary Fig. S5F-H**). Furthermore, *Pros1* and *Mertk*, a ligand-receptor pair that mediates immune suppression in human UM and that is upregulated by *BAP1* loss (23,24), is highly expressed in Neural Crest-like UM cells and tumor-associated macrophages (**Supplementary Fig. S5I,J**). These findings indicate that our Gnaq^CA^; Bap1^CKO^; Myc^CKI^ model of UM recapitulates key characteristics of human Class 2 UM, with the Neural Crest-like state particularly resembling the Class 2-like subtype.

### Key transcriptional programs underlying UM cell state diversity

To investigate the regulatory landscape of transcriptional programs across UM subpopulations, we determined the enrichment of transcription factor (TF) motifs among genes expressed in the different cell states. We identified the top 15 enriched TF motifs and visualized their activity across individual cells using a Z-score normalized heatmap (**Fig. 5A**). Hierarchical clustering of both TF motifs and cells revealed distinct expression modules that correspond to diverse transcriptional states. Motifs of KLF4, JUNB, EGR1, FOSB, YBX1, JUND, ATF4, ATF3, JUN, and MYC were highly enriched in cells of the Melanocytic subtype, while motifs of RBPJ, KLF6, RUNX, TEAD1, and FOXP2 were highly enriched in cells of the Neural Crest-like subtype (**Fig. 5A**). The enrichment of the MYC motif is in line with increased *Myc* expression in cells of the Melanocytic subtype (**Fig. 4A,B**). Notably, AP-1 family members, including FOSB, JUN, JUNB, and JUND, showed high activity predominantly in the Melanocytic subtype and, accordingly, their expression levels were higher in this subtype (**Fig. 5B-E**). Conversely, *Yap1* and *Tead1* are highly expressed in Neural Crest-like cells, which is consistent with the enrichment of the TEAD1 motif (**Fig. 5F, G**). Intriguingly, the expression of *Fosl1* strongly correlated with *Tead1* and *Runx1* rather than other AP-1 family members (**Fig. 5G-I**), and the gene product of *Fosl1*, FRA1, may form atypical complexes with TEAD1 to promote therapy resistance (25). Since AP-1 (via the MAPK pathway) and YAP are both critical downstream mediators of Gαq signaling, their respective enrichments in Melanocytic and Neural Crest-like subtypes highlights transcriptional heterogeneity in murine UM and underscores the involvement of distinct TF programs in defining UM cellular states and their lineage trajectories.

**Figure 5.**
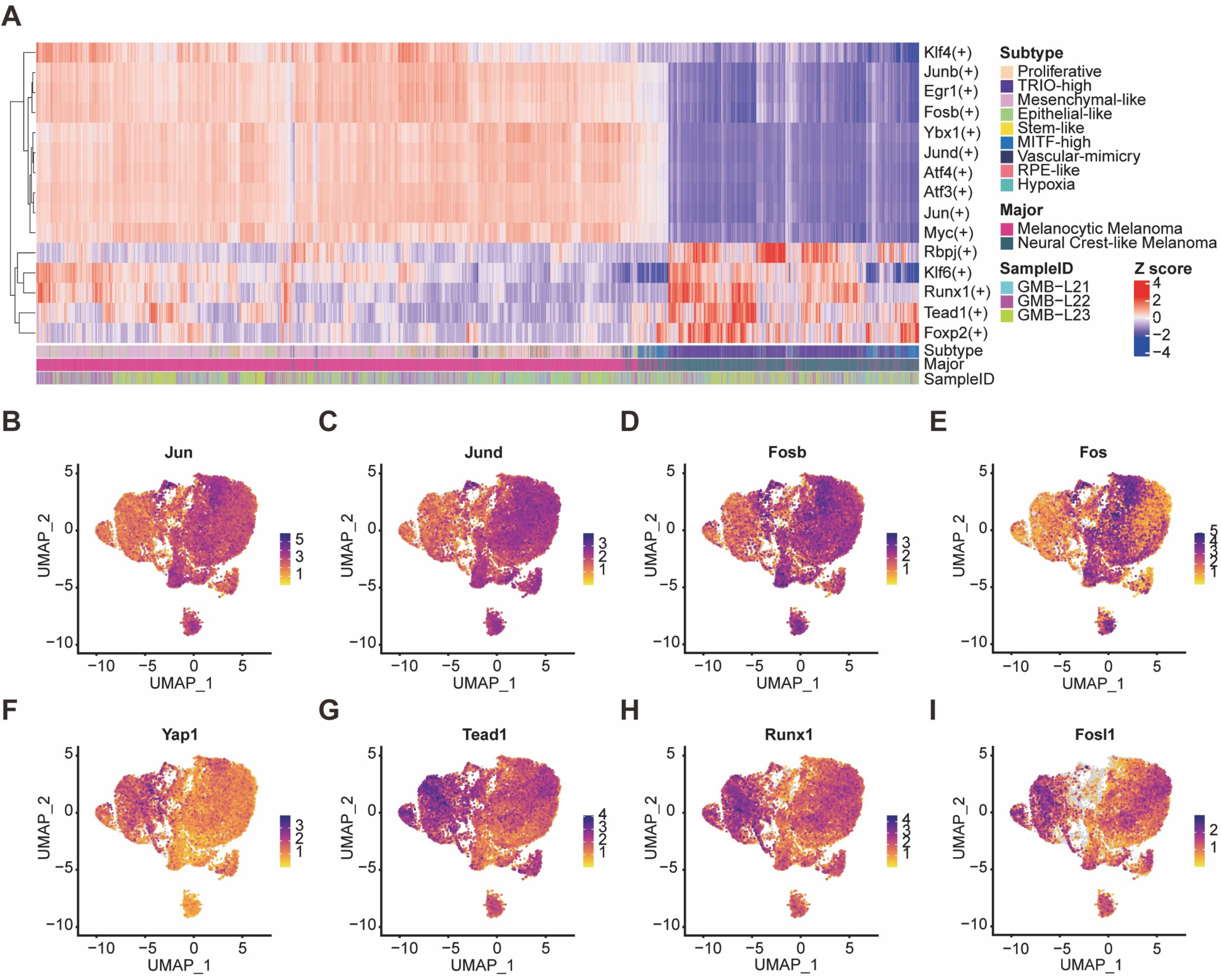
The transcription factor landscape across murine UM subtypes. **A,** Heatmap showing Z-score normalized transcription factor (TF) motif activity across mouse UM single cells from Gnaq^CA^; Bap1^CKO^; Myc^CKI^ tumors, as inferred by SCENIC analysis of the scRNA-seq data. Hierarchical clustering of both cells and motifs reveals distinct regulatory programs associated with Melanocytic and Neural Crest-like UM states. **B-I,** UMAP showing the expression of *Jun* (**B**), *Jund* (**C**), *Fosb* (**D**), *Fos* (**E**), *Yap1* (**F**), *Tead1* (**G**), *Runx1* (**H**), and *Fosl1* (**I**) in mouse UM cells from Gnaq^CA^; Bap1^CKO^; Myc^CKI^ tumors.

### Trajectory analysis reveals UM progression to a dedifferentiated state

To infer the dynamic transitions between transcriptional states within the malignant cell populations, we performed RNA velocity analysis using spliced and unspliced transcript abundances (**Fig. 6A**). Visualization of velocity vector fields projected onto the UMAP embedding revealed a strong directional flow from Melanocytic cells to Neural Crest-like cells (**Fig. 6A**). Two major paths were identified: the first path originated from the Mesenchymal-like cluster and arrived at the TRIO-high cluster via the MITF-high cluster; the other path originated from the Hypoxia cluster, passed through the Vascular-mimicry, Epithelial-like, and Stem-like clusters, and also ended at the TRIO-high cluster (**Fig. 6A**).

**Figure 6.**
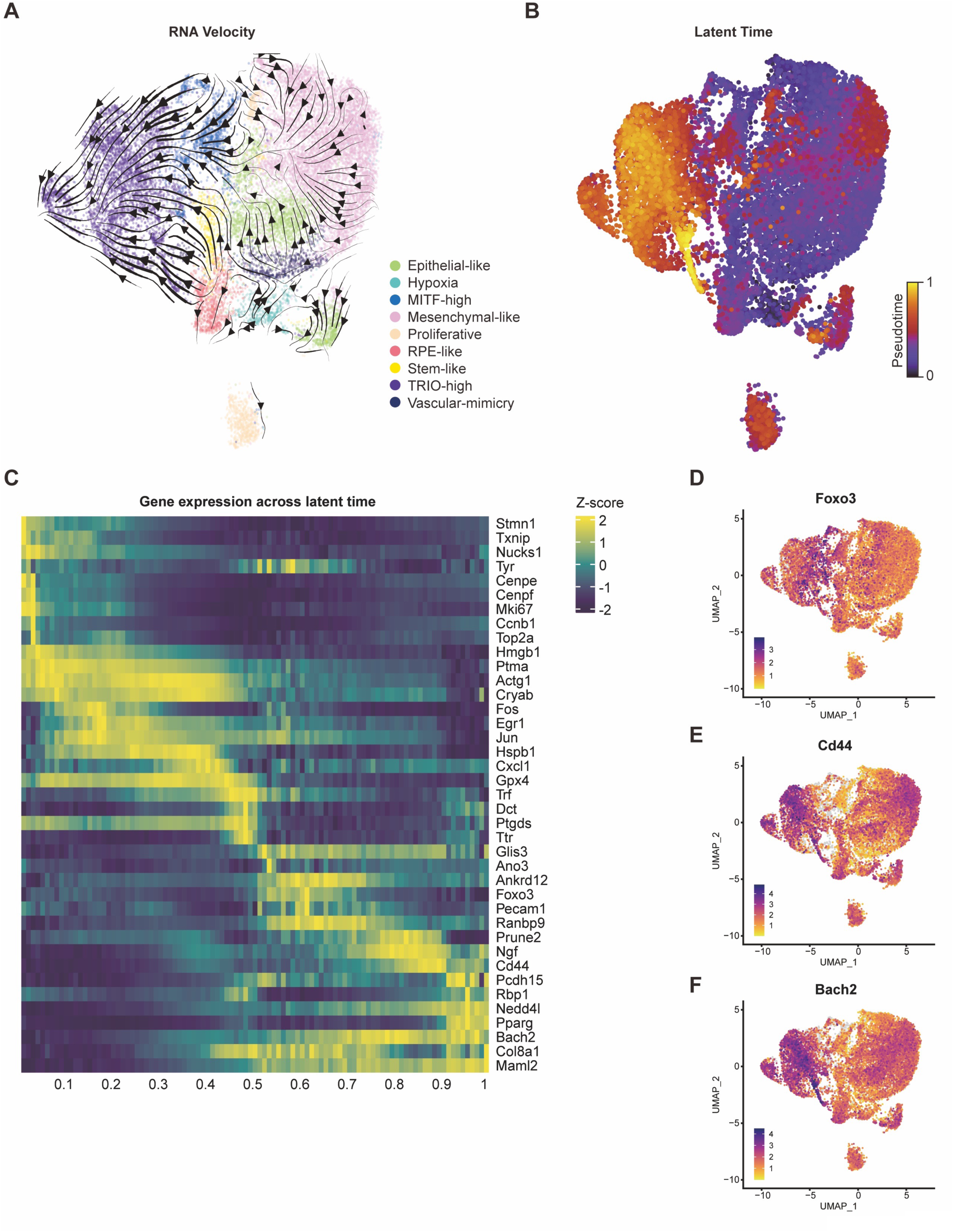
Progression trajectory analysis of murine UM. **A,** RNA velocity analysis based on spliced and unspliced transcript abundances revealed directional transitions across mouse UM cell states from Gnaq^CA^; Bap1^CKO^; Myc^CKI^ tumors. Velocity vector fields projected onto the UMAP embedding indicate major evolutionary paths. **B,** Latent time analysis shows the latency of single UM cells, color coded by pseudotime. **C,** Expression dynamics of representative genes plotted along latent pseudotime. **D-F,** UMAP plots showing the expression of *Foxo3* (**D**), *Cd44* (**E**), and *Bach2* (**F**) in murine UM cells.

To further reconstruct the continuous trajectory of transcriptional changes, we computed latent time, providing an unsupervised ordering of cells along inferred differentiation paths (**Fig. 6B**). Cells with the earliest latent time values predominantly resided within the Hypoxia cluster, and the Vascular-mimicry, Stem-like, and Epithelial-like clusters as well as parts of the Mesenchymal-like cluster were associated with early latent time as well (**Fig. 6B**). Intermediate latent times were associated with the Proliferative cluster and parts of the Mesenchymal-like cluster from the Melanocytic subtype, and the MITF-high cluster from Neural Crest-like subtype (**Fig. 6B**). The latest latent times were enriched in cells from the TRIO-high cluster. This latent time progression aligns with the RNA velocity analysis, suggesting a potential dedifferentiation trajectory from the Melanocytic to the Neural Crest-like subtype.

Analysis of gene expression dynamics across latent time further supported this model (**Fig. 6C, Supplementary Fig. S6**). Early in the latent time course, we observed high expression of cell proliferation and cell cycle genes, including *Cenpe*, *Cenpf*, *Mki67*, *Ccnb1*, and *Top2a* (**Fig. 6C, Supplementary Fig. S6**). As latent time progressed, cells upregulated the expression of immediate-early response genes (IEGs) such as *Fos*, *Egr1*, and *Jun* (**Fig. 6C, Supplementary Fig. S6**), indicating activation of stress-response pathways and transcriptional remodeling. At later stages of latent time, gene expression profiles shifted toward a more progenitor-like and migratory state, characterized by the induction of *Ngf*, *Cd44*, *Foxo3*, *Prune2* (**Fig. 6C-E, Supplementary Fig. S6**). These genes are associated with cell survival, motility, and stemness, hallmarks of neural crest-like properties. Additional late-expressed genes such as *Nedd4l*, *Pparg*, and *Bach2* further supported the emergence of a plastic, less differentiated state (**Fig. 6C, F, Supplementary Fig. S6**). Collectively, these gene expression dynamics are consistent with a model in which proliferative melanocytic UM cells undergo a progressive dedifferentiation process, potentially transitioning through stress-adaptive states toward a state with neural crest-like features.

### Genomic instability in murine UM correlates with transcriptional states and recapitulates copy number variation in human UM

To detect the copy number alteration (CNA) in murine UM, we applied inferCNV analysis to the scRNA-seq data using non-malignant immune cells as diploid reference. This analysis revealed two major CNA-defined tumor subpopulations, hereafter referred to as CNA Branch 1 and CNA Branch 2 (**Fig. 7A,B**). CNA Branch 1 exhibited broad chromosomal amplifications and deletions, while Branch 2 had a comparatively diploid profile, suggesting a spectrum of aneuploidy among tumor cells (**Fig. 7A)**. CNA Branch 1 was enriched for of Neural Crest-like cells, whereas Branch 2 was predominantly composed of Melanocytic cells (**Fig. 7B,C**). Functional state annotation further revealed that CNA Branch 1 was composed of almost all TRIO-high cells and approximately half of MITF-high cells, with minor contributions from Stem-like and Vascular-mimicry cells (**Fig. 7D**). In contrast, Branch 2 was predominantly comprised of cells from the seven minor subtypes that constitute the Melanocytic subtype (Vascular mimicry, Stem-like, Proliferative, Hypoxia, Epithelial-like, Mesenchymal-like, and RPE-like) (**Fig. 7D**). Overall, these data indicate a potential relationship between UM dedifferentiation and genomic instability.

**Figure 7.**
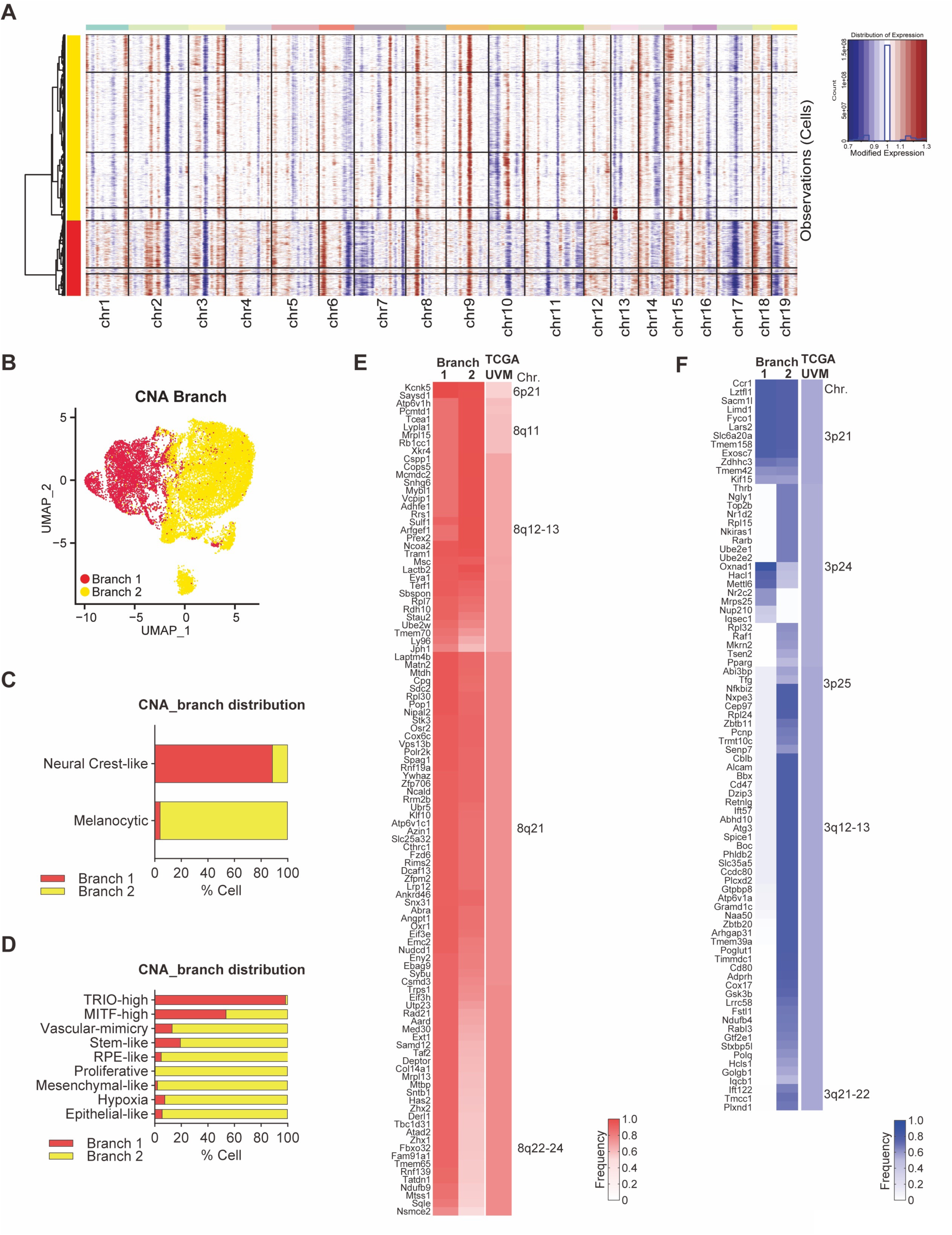
Copy number alterations correlate with UM subtypes. **A,** Inferred copy number alteration (CNA) profiles of individual malignant cells from murine UM tumors using inferCNV, using immune cells as diploid references. Heatmap shows relative genomic alteration levels across chromosomes, and hierarchical clustering of UM cells revealed two distinct CNA-defined UM branches. **B,** UMAP plot of mouse UM cells colored by CNA branch assignment. **C** and **D,** Proportional composition of CNA branch within each major subtypes (**C**) and phenotypic clusters (**D**). **E** and **F,** Heatmap of genes frequently amplified (**E**) or deleted (**F**) in both murine and human UM, color coded by frequency of chromosomal gain (**E**) or loss (**F**) in mouse UM cells or TCGA-UVM samples.

We observed a coordinated increase in gene expression across regions of mouse chromosomes 15, 1, and 5, syntenic to a stretch of human chromosome 8q (**Fig. 7A**). Additionally, regions of mouse chromosomes 3, 14, and 16, syntenic to part of human chromosome 3, displayed decreased expression and inferred CNA deletions (**Fig. 7A**). These CNAs potentially recapitulate chromosome 8q gain and loss of chromosome 3 in human Class 2 UM. By cross-species gene orthology mapping our CNA results to a uveal melanoma dataset in TCGA (TCGA-UVM), we identified 1,832 genes frequently altered in Branch 1 or Branch 2 cells (CNA in >50% murine UM cells) which are also gained or lost in human UM, of which 189 genes are frequently altered in human UM (CNA in >40% of human UM specimens). 105 genes with inferred copy number gains in murine UM are also frequently gained in human UM, with 2 genes on chromosome 6p and all others on chromosome 8q (**Fig. 7E**). 84 genes with inferred copy number losses in murine UM are also frequently lost in human UM, and all of them localize on chromosome 3 (**Fig. 7F**). Notably, while alterations of all 105 gained genes were present in both Branch 1 and Branch 2, only 12 of the 84 lost genes were shared by Branch 1 and Branch 2, indicating different patterns and trajectories of copy number gains and losses between the two branches. These findings indicate that our mouse model of UM displays CNA patterns that mimic some key genomic features observed in human UM, particularly patterns reminiscent of gain of chromosome 8q and loss of chromosome 3 seen in human Class 2 UM.

## DISCUSSION

Current UM mouse models, while establishing GNAQ/11 as initiating oncogenes, develop aggressive tumors within weeks and require euthanasia due to concurrent cutaneous disease (12,13), limiting their utility for studying immune interactions and metastatic progression. Our model addresses these limitations by providing (i) spatiotemporal control over oncogene activation restricted to the eye, (ii) stepwise genetic progression that models human disease evolution (26,27), (iii) immune competence enabling studies of tumor-immune interactions, and (iv) derivation of transplantable cell lines with distinct organ tropism.

Our model follows a genetic progression model for UM development, where GNAQ^Q209L^ activation induces choroidal nevi with limited penetrance, *BAP1* loss enhances nevus formation, and MYC activation promotes the formation of fully penetrant intraocular tumors resembling human UM. This stepwise progression models aspects of the multistage evolution observed in human UM, where *GNAQ/11* mutations are found in benign nevi, while BAP1 loss and chromosome 8q gains are associated with malignant progression and metastatic risk. While it remains unclear whether MYC is the key gene on 8q that drives the evolutionary pressure to gain more copies of 8q during human UM progression, our model provides experimental evidence that MYC expression, which may occur through chromosome 8q gains in human UM (11,17,27,28) can contribute to formation of intraocular tumors resembling UM in the mouse. Moreover, triple mutant Gnaq^CA^; Bap1^CKO^; Myc^CKI^ mice developed disseminated tumor cells in the liver, indicating that this model recapitulates not only primary tumor formation but also early steps in metastatic dissemination. Intravenous transplantation of primary UM cells led to metastasis in multiple distant organs, demonstrating the intrinsic metastatic potential of UM cells in this model.

Single-cell transcriptomic profiling revealed a complex and physiologically relevant tumor microenvironment composed of both malignant and non-malignant populations. Critically, this immune-competent models preserves the native tumor-immune ecosystem, with approximately 12% of tumors cells being immune infiltrates. The immune landscape of our model prominently featured *C1q*+ and *Arg1*+ macrophage subtypes as well as *Lag3*+ exhausted T cells and *Havcr2*+ dendritic cells (19,29–33), closely resembling the immunosuppressive microenvironment characteristic of human Class 2 UM. The presence of cancer-associated fibroblasts (including inflammatory, myofibroblast-like, and activated subtypes), combined with the diverse immune infiltrate, creates a tumor microenvironment that accurately reflects the complexity of human UM. This enables investigation of tumor-immune interactions and immunotherapies that are currently being explored clinically but lack adequate preclinical models for testing.

The malignant cells segregated into two major states: Melanocytic and Neural Crest-like. This phenotypic bifurcation recapitulates cellular hierarchies previously reported in cutaneous melanoma models (22), suggesting that despite differing anatomical locations and driver mutations, melanocytic malignancies may converge on common transcriptional programs during tumorigenesis. Sub-clustering further revealed phenotypic diversity, including proliferative, mesenchymal, epithelial-like, hypoxic, stem-like, and RPE-like states, reflecting the morphological heterogeneity observed in murine UMs and consistent with the known plasticity of melanoma cell states (34). The Melanocytic and Neural Crest-like cell populations exhibited distinct features, indicating different functional roles in tumor progression. Melanocytic cells showed high expression of MYC target genes and downstream metabolic, ribosomal, and mitochondrial programs, consistent with MYC coordinating cellular growth with anabolic metabolism and protein synthesis (35). The robust correlation between *MYC* and ribosomal gene expression observed in both our model and the TCGA-UVM dataset underscores a conserved regulatory axis in UM pathobiology. In contrast, Neural Crest-like cells exhibited lower MYC target expression and upregulation of genes associated with immune evasion (e.g., *Pros1*, *Mertk*) (23,24) and key clinical biomarkers of aggressive Class 2 UM, including *Htr2b*, *Ecm1*, and *Rab31* (10,36). Intriguingly, while *MYC* is upregulated in human Class 2 compared to Class 1 UMs, the *MYC* expression pattern in Melanocytic and Neural Crest-like subpopulations mirrors the findings in our mouse model, potentially indicating that *MYC* downregulation is associated with UM progression. This is supported by the clinical observation that high MYC expression levels correlate with better survival outcomes in UM patients (37), suggesting a nuanced role of MYC in UM progression.

Transcriptional network analysis revealed further divergence between these states. Melanocytic cells showed enrichment of MYC and AP-1 binding motifs, aligning with elevated expression of proliferation-related and metabolic genes. The AP-1 complex, which has been implicated in melanocyte lineage maintenance and melanoma progression (26,38), is a canonical effector of MAPK signaling and activated downstream of mutant GNAQ suggesting these cells are transcriptionally poised for proliferation and biosynthesis Neural Crest-like cells exhibited enrichment of motifs associated with YAP-TEAD signaling, RUNX1, FOXP2, RBPJ, and KLF6, corresponding with elevated expression of Yap1 and Tead1. This is notable as YAP is a key downstream target of GNAQ/GNA11 signaling (39,40), has been shown to drive dedifferentiation and stem-like properties (41,42), and may thus play a potential role in promoting cellular plasticity and metastatic competence in our model. The co-expression and motif co-enrichment of FRA1 (FOSL1) with TEAD1 is particularly intriguing, as their non-canonical interaction has been implicated in drug resistance (25).

The phenotypic segregation paralleled distinct patterns of genomic instability. Neural Crest-like cells showed broad chromosomal amplifications and deletions, suggesting dedifferentiation is associated with increased genomic instability. In contrast, Melanocytic cells displayed a more diploid-like profile, reflecting a genomically more stable phenotype. Cross-species mapping identified a substantial overlap between copy number alterations in murine and human UM, particularly gains of chromosome 8q and losses of chromosome 3 (43,44). Among the ∼600 protein-coding genes on 8q, 105 orthologous genes underwent copy number gains in our mouse model—an unexpectedly high concordance that underscores the fidelity of this model in recapitulating human UM genomics. Importantly, this broad amplification pattern suggests that multiple drivers beyond MYC are critical for UM progression. While MYC is a well-established oncogene on 8q, the coordinated amplification of numerous genes in the 8q11-13 and 8q21-24 regions indicates that UM progression likely depends on the cooperative effects of multiple 8q-resident genes rather than MYC alone. The fact that our model recapitulates this broad 8q amplification pattern suggests that the selective pressure for 8q gain in UM reflects the need for multiple cooperating oncogenes in this chromosomal region. Deletions corresponding to human chromosome 3 varied between Melanocytic and Neural Crest-like cells, suggesting distinct evolutionary trajectories or selective pressures in these subtypes. The biological significance of this divergence remains unclear but may reflect differential lineage constraints within each subtype. The distinction in CNA patterns further suggests that genetic instability may drive cell state transitions in UM, providing a mechanistic link between dedifferentiation, chromosomal aberrations, and tumor progression.

Our trajectory analysis suggested a directional transition from the Melanocytic to the Neural Crest-like state, accompanied by dynamic regulation of key transcriptional programs. Dedifferentiation toward a neural crest-like state has also been observed in cutaneous melanoma and is associated with immune evasion, metastasis, and resistance to therapy (22,45–47). The induction of immediate early genes (*Fos*, *Egr1*, *Jun*) at intermediate stages suggests transcriptional adaptation to cellular stress (48) while the late expression of stem cell-associated genes (*Foxo3*, *Cd44*, *Ngf*, *Prune2*) points to acquisition of stem-like and migratory traits—features commonly attributed to invasive or therapy-resistant cells (49,50). However, without definitive lineage tracing, we cannot exclude the alternative possibilities that these populations diverge from a common ancestor early during tumor evolution or arise independently from distinct cells of origin.

In summary, our study introduces a biologically and preclinically relevant mouse model of human UM that recapitulates many key features of the human disease, including cell state heterogeneity, genomic alterations, and metastatic potential. Our work demonstrates that overexpression of MYC can substantially increase the malignant potential of these mouse tumors, although this remains to be verified in human UM. Interestingly MYC can cooperate with *GNAQ* mutation and *BAP1* loss to enhance malignant transformation and its spontaneous downregulation by tumor cells is associated with a transition to the Neural Crest-like state. The identification of distinct Neural Crest-like and Melanocytic subtypes, each with unique transcriptional signatures and copy number landscapes, adds to a growing appreciation of UM as a heterogeneous and dynamic disease, as also highlighted by scRNA-seq analyses of human UM (19). This immune-competent model provides a powerful platform for advancing UM research by enabling studies of drivers of metastatic progression such as PRAME (51) and GDF15 (52), tumor-immune interactions, and preclinical testing of immunotherapies – critical needs given the limited treatment options for metastatic disease.

### Limitations of this study

While our mouse model represents a significant advance in UM research, we acknowledge certain limitations. The lentiviral Cre delivery approach may potentially result in tumors with atypical cellular heterogeneity, as suggested by the RPE-like and epithelial-like malignant populations identified in our scRNA-seq analysis. This heterogeneity could reflect the biology of human UM, which also displays phenotypic plasticity, but might also result from non-specific recombination events in different ocular cell types such as RPE cells. Despite this limitation, our model successfully recapitulates molecular signatures and phenotypic states of human UM, particularly the aggressive Class 2 subtype. Future iterations of the model will incorporate lineage-specific Cre drivers to improve resolution in defining the initiating cell population.

## MATERIALS AND METHODS

### Generation of a GNAQ^Q209L^ (Gnaq^CA^) allele

A 9,217bp genomic fragment from the murine *Gnaq* locus, including 4,491bp upstream and 4,596bp downstream of exon 5, were inserted into pKO2.2 (Addgene plasmid #22676) by InFusion cloning. Exon 5 was then mutated at nucleotide 626 (AàT) of the *Gnaq* coding sequence by site-directed mutagenesis to create the Q209L hotspot mutation. A minigene cassette was generated by GeneArt (ThermoFisher) and inserted into an NdeI site 649bp upstream of exon 5. This minigene cassette contained the following elements: (i) a Splice Acceptor followed by the partial Gnaq cDNA from exon 5 to exon 7 and two SV40 polyA signals; (ii) a PGK promoter followed by the Blasticidin resistance marker and a BGH polyA signal in the reverse orientation; (iii) FRT sites flanking the PGK promoter and the BGH polyA to enable Flp recombinase-mediated removal of the Blasticidin selection cassette; and (iv) loxP sites flanking the upstream Splice acceptor and the downstream FRT site to enable Cre recombinase-mediated removal of the entire minigene cassette. The final targeting vector was sequence verified by Plasmidsaurus whole plasmid sequencing. V6.5-C10 embryonic stem cells (53) on a mixed C57BL/6 x 129S4/SvJae background were targeted by homologous recombination in Moffitt’s Gene Targeting Core and selected in 10 µg/ml Blasticidin. Clones were screened by Southern blot using a 5’ external probe (NheI digest), a 3’ external probe (AflII digest), and a Blasticidin internal probe (HindIII digest). The presence of the Q209L mutation was confirmed by Sanger sequencing in Southern blot-positive clones. Clones were injected into Balb/c blastocysts and transferred into pseudopregnant CD1 females. High contribution chimeras were then bred to C57BL/6 females to achieve germline transmission.

#### Animals

To generate a genetically engineered mouse strain harboring conditional alleles for *Gnaq*, *Cas9*, and *Myc*, we performed a multi-generational breeding scheme using the following strains: GNAQ^Q209L^ (Gnaq^CA^) mice were generated in this study as described above, Rosa26-LSL-Cas9-EGFP (Cas9^CKI^) mice were purchased from The Jackson Laboratory (#028551), H11-LSL-Myc^T58A^-Luciferase (Myc^CKI^) mice were obtained from Elsa Flores. Gnaq^CA^ were maintained on a mixed C57BL/6 x 129S4/SvJae background. Cas9^CKI^ and Myc^CKI^ strains were maintained on a C57BL/6J background. Initial crosses were performed to generate double homozygous Gnaq^CA^; Cas9^CKI^ breeders, which were then intercrossed to homozygous Myc^CKI^ mice to obtain the desired triple mutant animals. Genotyping was performed on genomic DNA extracted from tail biopsies using PCR with allele-specific primers (listed in **Supplementary Table S1**). Only mice carrying the correct allele combinations were used for experiments. All procedures involving animals were approved by the Institutional Animal Care and Use Committee (IACUC) and conducted in accordance with institutional and national guidelines.

#### Mouse embryonic fibroblasts

Mouse embryonic fibroblasts (MEFs) were isolated from Gnaq^CA^ embryos at E13.5 and infected with lentiviral Cre. RNA was extracted from MEFs using TRIzol (Invitrogen) following protocols supplied by the manufacturer. cDNA was generated with PrimeScript RT Master Mix (Takara). *Gnaq* exon 5 was PCR amplified with allele-specific primers (listed in **Supplementary Table S1**) using GoTaq Green Master Mix (Promega) and subjected to Sanger sequencing. MEFs isolated from Gnaq^CA^; Cas9^CKI^ mice were infected with pL-CLB. Genomic DNA (gDNA) was isolated from MEFs using a standard Proteinase K digestion and ethanol precipitation method. *Bap1* exons 1-3 were amplified by PCR using primers listed in **Supplementary Table S1** from gDNA with GoTaq Green Master Mix (Promega) and subjected to Sanger sequencing.

#### Lentivirus production, concentration, and suprachoroidal administration

pl-CMV-Cre-EF1a-luciferase-U6-sgBap1 (pL-CLB), which expresses Cre recombinase from the CMV promoter, firefly luciferase from the EF1α promoter, and a U6-driven sgRNA targeting Bap1, was generated based on a pLenti construct. The sequence of the sgRNA is listed in **Supplementary Table S1**. To produce lentivirus supernatants, HEK293T cells were transfected with lentiviral vector and delta8.2 and VSVG helper plasmids at a 9:8:1 ratio using JetPRIME transfection reagent. Virus-containing supernatant was collected at 48 and 72 hours post-transfection, filtered through a 0.45 μm PVDF filter, and stored at 4°C. Supernatants were concentrated using Lenti-X Concentrator (Takara Bio): viral supernatant was mixed with 1/3 volume of Lenti-X reagent, incubated at 4°C overnight, and centrifuged at 1,500 × g for 45 minutes at 4°C. Pellets were resuspended in sterile PBS (50× concentration) and aliquots stored at –80°C. Viral titer was quantified using the Lenti-X qRT-PCR Titration Kit.

Suprachoroidal administration of concentrated lentivirus was performed in adult mice (6–8 weeks old) under isoflurane anesthesia. A 33-gauge beveled Hamilton syringe was inserted tangentially through the sclera to access the suprachoroidal space. 2  μL of concentrated lentivirus (titer ∼10⁷–10⁸ TU/mL) was slowly injected over 30-60 seconds. To prevent reflux, the needle was held in place for 30 seconds post-injection before removal. The eye was treated with topical erythromycin ophthalmic ointment post-procedure. Efficient gene delivery was verified by bioluminescence imaging (IVIS) at 3–5 days post-injection.

#### Orthotopic and metastatic transplant of murine uveal melanoma

Uveal melanomas were harvested from *Gnaq^CA^; Bap1^CKO^; Myc^CKI^* mice when ocular enlargement was visibly apparent. Mice were euthanized, and eyes were enucleated using fine scissors and forceps under a dissection microscope. After removal, each eye was placed in ice-cold sterile PBS. Under a stereomicroscope, extraocular tissues (conjunctiva, optic nerve), lens, and vitreous were removed. Tumor tissue arising from the choroid and adjacent structures was dissected out with fine-tipped forceps and immediately transferred into the enzyme solution. Enzymatic dissociation was performed using the Tumor Dissociation Kit (Miltenyi Biotec) according to the manufacturer’s instructions. Tissues were incubated at 37°C for 30–45 minutes with intermittent gentle agitation. The resulting cell suspension was washed with PBS containing 2% FBS and centrifuged at 300 × g for 5 minutes. Cells were resuspended in sterile PBS, counted using a hemocytometer, and assessed for viability by trypan blue exclusion. All downstream injections were performed immediately following dissociation to minimize ex vivo artifacts.

Orthotopic tumor engraftments were performed in 6–8-week-old NSG mice under isoflurane anesthesia. A 33-gauge beveled Hamilton syringe was inserted tangentially through the sclera to access the suprachoroidal space. 20,000 freshly dissociated uveal melanoma cells in 2 µL of sterile PBS were slowly injected over 30-60 seconds. To prevent reflux, the needle was held in place for 30 seconds post-injection before removal. The eye was treated with topical erythromycin ophthalmic ointment post-procedure. Tumor development was tracked by in vivo imaging, and mouse eyes bearing uveal melanoma tumors were collected at defined time points for histologic assessment.

To assess metastatic potential, 1 million freshly isolated uveal melanoma cells in 100 µL sterile PBS were injected into the lateral tail vein of anesthetized NSG mice using a 30-gauge insulin syringe. Metastasis development was tracked by in vivo imaging and histologic assessment at defined time points. Liver, lung, and other organs were harvested at endpoint for histologic evaluation of metastases.

#### Immunohistochemistry and Periodic Acid–Schiff (PAS) Staining

Mouse eyes bearing UMs and organs bearing metastasis were harvested and fixed in 10% neutral-buffered formalin for 48-96 hours at room temperature. Embedding, sectioning, and H&E staining were performed by IDEXX BioAnalytics.

For standard Immunohistochemistry (IHC), slides were deparaffinized in xylene and rehydrated through graded ethanol. Antigen retrieval was performed using citrate buffer (pH 6.0) in a microwave oven for 12 minutes. Endogenous peroxidase activity was quenched using 3% hydrogen peroxide for 10 minutes. Sections were blocked in 2.5% normal horse serum for 1 hour at room temperature and incubated overnight at 4°C with primary antibodies: anti-Tyrosinase (1:100, Thermo Fisher, PA5-86066) and anti-Melan-A (1:100, Thermo Fisher, MA5-13232). Detection was performed using HRP-conjugated secondary antibodies (Vector Labs) and DAB substrate (Vector DAB Peroxidase Substrate Kit), followed by hematoxylin counterstaining. Slides were dehydrated and mounted with permanent mounting media.

Fluorescent Immunohistochemistry (F-IHC) was performed using antibodies against RPE65 (1:150, GeneTex, GTX13826) and SOX10 (1:50, Cell Signaling Technology, 78330). After antigen retrieval and blocking as above, sections were incubated overnight at 4°C with primary antibodies diluted in 1% BSA in PBS. The next day, slides were washed and incubated with species-appropriate Alexa Fluor-conjugated secondary antibodies (1:1,000, Thermo Fisher) for 1 hour at room temperature. Slides were coverslipped using ProLong Gold Antifade Mountant with DAPI (Thermo Fisher) and imaged using a fluorescence microscope (EVOS-Auto).

PAS staining was performed using the Periodic Acid–Schiff Stain Kit (Sigma-Aldrich) according to the manufacturer’s instructions. Briefly, paraffin sections were deparaffinized and rehydrated, then oxidized in 0.5% periodic acid for 5 minutes, rinsed, and incubated in Schiff’s reagent for 15 minutes. Sections were counterstained with hematoxylin, dehydrated, and mounted. PAS-positive structures were visualized as magenta.

#### Western blot analysis

Fifteen micrograms of protein were separated on NuPAGE 4% to 12% precast gels (Thermo Fisher Scientific) and transferred to nitrocellulose membranes. Membranes were blocked in 5% non-fat dry milk in TBST and incubated with one of the following primary antibodies overnight at 4°C: Gαq (1:1000, ,14373), ERK1/2 (1:2,000, Cell Signaling Technology, 4695), p-ERK (Thr202/Tyr204) (1:1,000, Cell Signaling Technology, 4370), YAP (1:1,000, Cell Signaling Technology, 4912), p-YAP (Ser127) (1:1,000, Cell Signaling Technology, 4911), BAP1 (1:1,000, Abcam, ab255611). Anti-beta-actin (1:3,000, Invitrogen, AM4302) was blotted as a loading control. Membranes were washed 3 times with TBST for 10 min, followed by incubation with HRP-conjugated secondary antibodies (1:3,000) for 1 hour at room temperature. After three washes in TBST, chemiluminescence substrate (1:1) was applied to the blot for 4 min and chemiluminescence signal was captured using a LI-COR imaging system.

#### Single-cell sequencing

Single-cell RNA-sequencing was performed using the 10X Genomics Chromium System (10X Genomics) by Moffitt’s Molecular Genomics Core. Cell viability and counts were obtained by AO/PI dual fluorescent staining and visualization on the Nexcelom Cellometer K2 (Nexcelom Bioscience LLC). Cells were then loaded onto the 10X Genomics Chromium Single Cell Controller to encapsulate approximately 10,000 cells per sample. Single cells, reagents, and 10X Genomics gel beads were encapsulated into individual nanoliter-sized Gelbeads in Emulsion (GEMs), and reverse transcription of polyadenylated mRNA was performed in each droplet at 53°C. The cDNA libraries were then completed in a single bulk reaction by following the 10X Genomics Chromium GEM-X Single Cell 3’ Reagent Kit v4 user guide, and 50,000 sequencing reads per cell were generated on the Illumina NovaSeq6000 instrument. Demultiplexing, barcode processing, alignment, and gene counting were performed using the 10X Genomics CellRanger v8 software.

#### scRNA-seq data processing, filtering, batch effect correction, and clustering

Raw sequencing reads from scRNA-seq were processed using Cell Ranger (v7.1.0, 10X Genomics). Briefly, the base call (BCL) files generated by Illumina sequencers were demultiplexed into fastq files based on the sequences of the sample index, and aligned against GRCm39 mouse transcriptome using STAR (54). Cell barcodes and UMIs associated with the aligned reads were subjected to correction and filtering. Filtered gene-barcodes matrices containing only barcodes with UMI counts passing the threshold for cell detection were imported to Seurat v4.0 (55) for downstream analysis. Barcodes with fewer than 200 genes expressed or more than 10% UMIs originated from mitochondrial genes were filtered out; genes expressed in fewer than 3 barcodes were also excluded. This process resulted in 24,455 cells from three mice. For each sample, standard library size and log-normalization were performed on raw UMI counts using *NormalizeData*, and the top 5,000 most variable genes were identified by the “vst” method in *FindVariableFeatures*.

Individual samples were further integrated to remove batch effects using an anchor-based method (56) implemented in Seurat v4.0 using *FindIntegrationAnchors* and *IntegrateData* functions in Seurat with 8,000 “anchors” and top 40 principal components. Briefly, dimension reduction was performed on each data using diagonalized canonical correlation analysis (CCA) and L2-normalization was applied to the canonical correlation vectors to project the datasets into a shared space. The algorithms then searched for mutual nearest neighbors (MNS) across cells from different datasets to serve as “anchors” which encoded the cellular relationship between datasets. Finally, correction vectors were calculated from “anchors” and used to integrate datasets.

From the integrated data, scaled z-scores for each gene were calculated using *ScaleData* function in Seurat by regressing against the percentage of UMIs originated from mitochondrial genes, S and G2/M phases scores, and total reads count. A shared nearest neighbor (SNN) graph was constructed based on the first 40 principal components computed from the scaled integrated data. Louvain clustering was performed using the *FindClusters* function at resolution 1.2 for major cell type scRNA-seq data (34 clusters). Uniform manifold approximation and projection (UMAP) was used to visualize single-cell gene expression profile and clustering, using *RunUMAP* function in Seurat with default settings. Differential expression analysis was performed using *FindAllMarkers* function in Seurat with logfc.threshold=0.25, min.pct=0.2, and test.use=”wilcox”. Cells within each cluster were compared against all other cells. Genes with Bonferroni-corrected p-value < 0.05 and an average log-fold change > 0.25 were considered differentially expressed. Clusters were annotated by comparing differential genes with canonical markers for major populations: Melanoma cells (neural crest-like features: *Mitf, Mlana, Tyr, Gli2, Meis1, Sox5*; melanocytic features: *Mitf, Mlana, Tyr, Dct, Tyrp1)*, B cells (*Cd79a, Cd79b, Cd22*), T/NK cells (*Cd3d*, *Cd3*e, *Cd3*g), Monocytes (*Ly6c2, Vcan, Plac8*), Neutrophils (*S100a8, S100a9*), mregDC (CCr7, *Fscn1, Ccl22, Cacnb3*), Multiciliated epithelial cells (*Crocc2*, *Foxj1, Deup1, Mcidas*), Endothelial cells (*Pecam1, Cdh5, Tie1*), Inflammatory CAF (*Col1a1*, *Col1a2*, *Col5a1*, *Cxcl5*, *Il11*, *Saa3*), Myofibroblast-like CAF (*Col1a1*, *Col1a2*, *Col5a1*, *Actg2*, *Cd248*, *Myh11*), Activated CAF (*Col1a1*, *Col1a2*, *Col5a1*, *Adamts2*, *Col6a3*, *Tnc*) , Arg1+ Macrophages (*Cd68, Mrc1, Arg1, Ccl9, Frt3*), C1qa+ Macrophages (*Cd68, Mrc1, C1qa, C1qb, C1qc, Fcrls*), RPE cells (*Rbp1, Rlbp1, Ttr*), Retinal bipolar cells (*Isl1, Neurod4, Pcp2*), Retinal photoreceptor cells (*Gnat1, Gngt1, Rho*), and Müller glial cells (*Aqp4, Gtap, Sox2*).

#### Annotation of melanoma cells

Melanoma cells were annotated as Neural Crest-like (*Gli2, Glis3 Meis1, Sox5*) and Melanocytic (*Mlana, Dct, Tyrp1, Slc24a5*) and these two major populations were further annotated into nine subtypes. Neural Crest-like melanoma cells: TRIO-high (*Trio, Ngf, Camta1, Arhgap10*) and MITF-high (*Tcf4, Nfia, Mift, Gsk3b*); Melanocytic melanoma cells: Epithelial-like (*Prdx1, Krt18, Gpnmb, Atp1b1*), Mesenchymal-like (*Vim, Pcolce, Mfap4, Acta2*), Proliferative (*Top2a, Stmn1, Pcna, Mki67*), Hypoxia (*Ndufa4l2, Higd1a, Egln3, Bnip3*), Stem-like (*Pdgfc, Igfbp2, Gdf15, Gas6*), Vascular-mimicry (*Scin, Prtg, Pecam1, Itgb6*), and RPE-like (*Ttr, Rpe65, Rlbp1, Rdh5, Rbp1*). Differential expression analysis was performed to compare two major populations of melanoma cells, as well as the 9 subtypes, using *FindMarkers* function in Seurat with logfc.threshold=0.25, min.pct=0.2, and test.use=”wilcox”. Genes were ranked based on -log10(p-value)*(sign of log2(fold-change)). Pre-ranked GSEA was performed on gene rankings using R package fgsea (57) with 10,000 permutations, against Hallmark, REACTOME, and GO databases from MsigDB (58–60). The normalized enrichment scores (NES) of gene sets were visualized using heatmaps. To further characterize the malignant cells, we compared gene expression of 9 subtypes to cell identity signatures previously reported in mouse cutaneous melanoma studies. Enrichment scores of these signatures were calculated for each malignant cell using the *AUCell* algorithm implemented in SCENIC (61), and the scores were visualized in heatmaps.

Copy number variation patterns in malignant cells were extracted using InferCNV (62) R package v1.20.0. Normal immune cells identified above (B, T/NK, Neutrophils, mregDC and Monocytes) were selected as “reference” cells for de-noise control. InferCNV analysis was performed using “denoise” mode to correct for batch effects from different mice, with tumor_subcluster_partition_method = ‘qnorm’, HMM=TRUE, and analysis_mode = ’subclusters’. The “cluster by group” parameter was turned off to allow the observation cells to cluster unbiasedly based on CNA patterns. Two major CNA branches were identified in melanoma cells, corresponding to the two main melanoma cell types: neural crest-like melanoma cells and melanocytic melanoma cells.

To compare CNV patterns between mouse and human UM, we mapped CNV results obtained from mouse UM cells as described above to the TCGA uveal melanoma dataset (TCGA-UVM). CNV data for TCGA-UVM were downloaded from cBioPortal. *HOMDEL* and *HETLOSS* events were defined as loss, and *GAIN* and *AMP* events were defined as gain. Mouse genes were mapped to their human orthologs using the Mouse Genome Informatics database (www.informatics.jax.org). Genomic locations of human orthologs were retrieved using the *biomaRt* package in R. The CNV status of these human orthologs was then summarized in the TCGA-UVM dataset to evaluate conservation of CNV patterns between species.

RNA velocity analysis was performed to infer the dynamic states of malignant cells. Initially, spliced and unspliced transcript abundances were quantified from BAM files generated from CellRanger count using Velocyto (63). The resulting loom files were merged across samples. Seurat meta data was integrated with the loom file and malignant cells were extracted for downstream analysis. The dynamical model of RNA velocity in Python module scVelo (64) was applied to estimate transcriptional rates and compute velocity vectors for individual cells in “dynamical” mode. Briefly, we used scv.tl.recover_dynamics() to infer gene-specific transcriptional kinetics from spliced and unspliced mRNA counts. Velocity vectors were computed using scv.tl.velocity() in dynamical mode, followed by scv.tl.velocity_graph() to estimate the transition probabilities between cells. Latent time—a continuous pseudotime derived from the velocity field—was computed with scv.tl.latent_time(), representing the internal clock of cells along their differentiation trajectories. The velocity fields were visualized on the batch-corrected UMAP projection, with arrows overlaid to indicate the direction and magnitude of transcriptional changes. To identify genes associated with the latent time progression, malignant cells were grouped into 100 bins based on their estimated latent time. Marker genes specific to each bin were identified using the *FindAllMarker* function in Seurat and visualized in a heatmap.

#### SCENIC analysis

Transcription factor motif analysis was performed using pySCENIC (61) on a normalized expression matrix of the 18,901 melanoma cells with default setting. In addition, option --mask_dropouts was used. Using the mouse v10 database (https://resources.aertslab.org/cistarget/databases/mus_musculus/mm10/; https://resources.aertslab.org/cistarget/motif2tf/), 323 transcription factors were identified in 18,901 melanoma cells. The resulting matrix was z-scored using the *scale* function from base R. Transcription factors with the top 15 highest AUC scores were visualized in a heatmap with hierarchical clustering.

#### Myc-luciferase transgene sequence search

We performed a BLAT search between the Firefly Luciferase sequence and 90 bp 10X raw reads from all three samples. 10X reads that matched the *Firefly* Luciferase sequence were defined as those with 100% identity and an alignment length of at least 89 bp. Cells containing “*FireflyLuc* reads” were extracted and counted how many such reads were associated with each unique cell ID. The presence of any FireflyLuc reads in a given cell ID was recorded as “FireflyLuc Yes”; absence was marked as “FireflyLuc No”. The number of FireflyLuc reads per unique cell ID was recorded as LucReadNum, which was used to represent the expression level of the Myc^T58A^-IRES-Luciferase transgene in each cell. We compared gene expression of the 18,901 melanoma cells to the mouse MYC targets signatures in Hallmarks. Enrichment scores of these signatures were calculated for each malignant cell using the *AUCell* algorithm implemented in SCENIC (61), and the scores were visualized in heatmaps. All this information was visualized by Seurat v4.0 using *FeaturePlot*. Spearman correlation was performed between *Myc* and other genes to identify genes that have the highest correlation with *Myc* expression.

## SUPPLEMENTARY FIGURE LEGENDS

**Figure S1.**
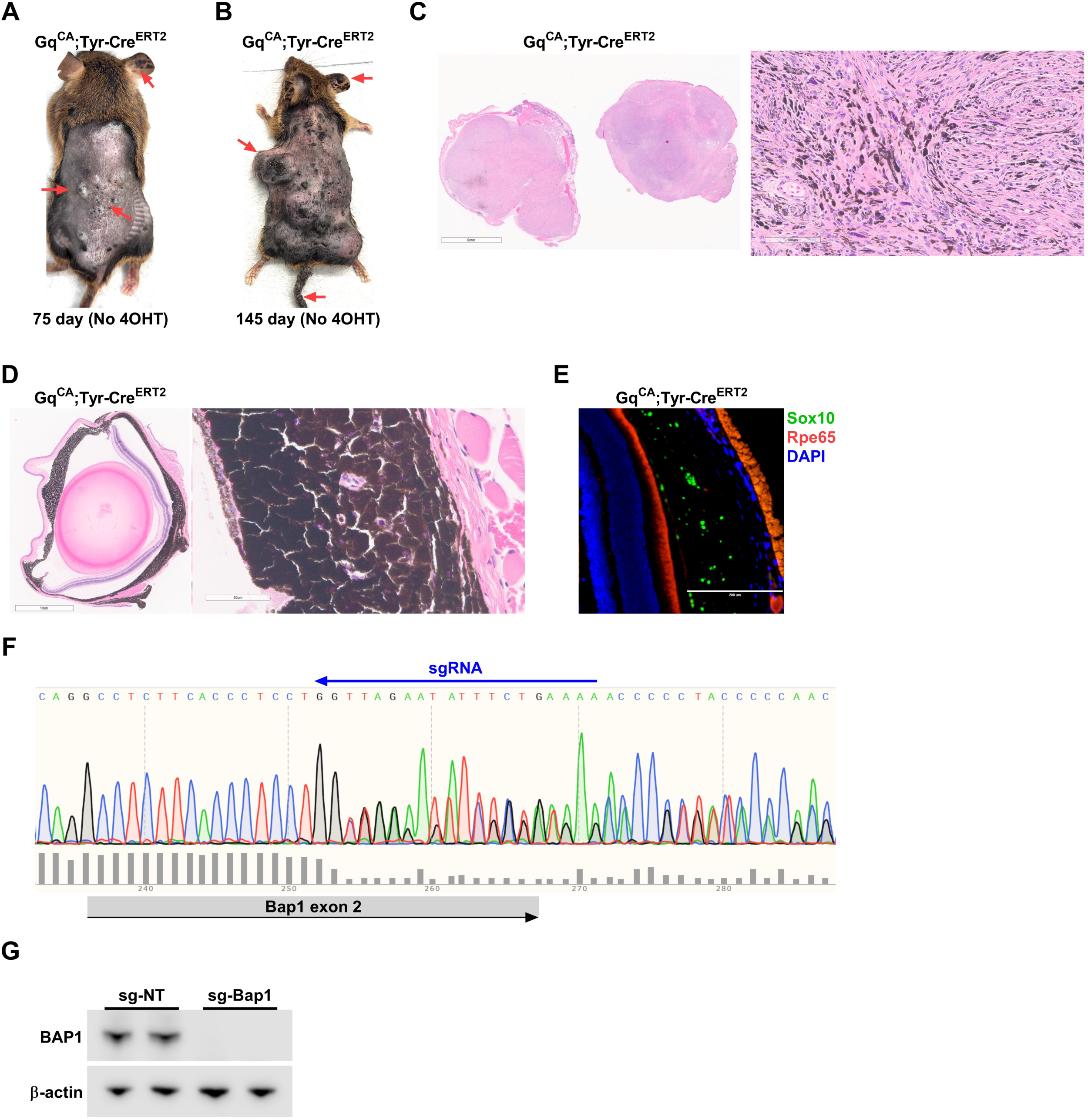
Phenotype of Gnaq^CA^; Tyr-Cre^ERT2^ mice and validation of Bap1 deletion. **A** and **B,** Representative images of Gnaq^CA^; Tyr-Cre^ERT2^ mice showing cutaneous nevi and small tumors at 75 days of age (**A**) and extensive cutaneous melanomas at age of 145 days (**B**). Melanomas formed in the absence of 4-Hydroxytamoxifen (4OHT) administration, indicating spontaneous Cre recombination. **C,** Representative images of H&E staining of skin tumors from Gnaq^CA^; Tyr-Cre^ERT2^ mice. **D,** Representative images of H&E staining of eyes of Gq^CA^; Tyr-Cre^ERT2^ mice show extensive hyperplasia in choroid and iris. **E,** Representative fluorescent IHC showing SOX10 and RPE65 expression in uveal tracts of Gnaq^CA^; Tyr-Cre^ERT2^ mice. **F,** Mouse embryonic fibroblasts (MEFs) were isolated from Gnaq^CA^; Cas9^CKI^ mice and infected with pL-CLB. PCR amplification of genomic DNA followed by Sanger sequencing confirmed CRISPR knockout of Bap1. **G,** Western blot showing loss of BAP1 expression in Gnaq^CA^; Cas9^CKI^ MEFs infected with pL-CLB.

**Figure S2.**
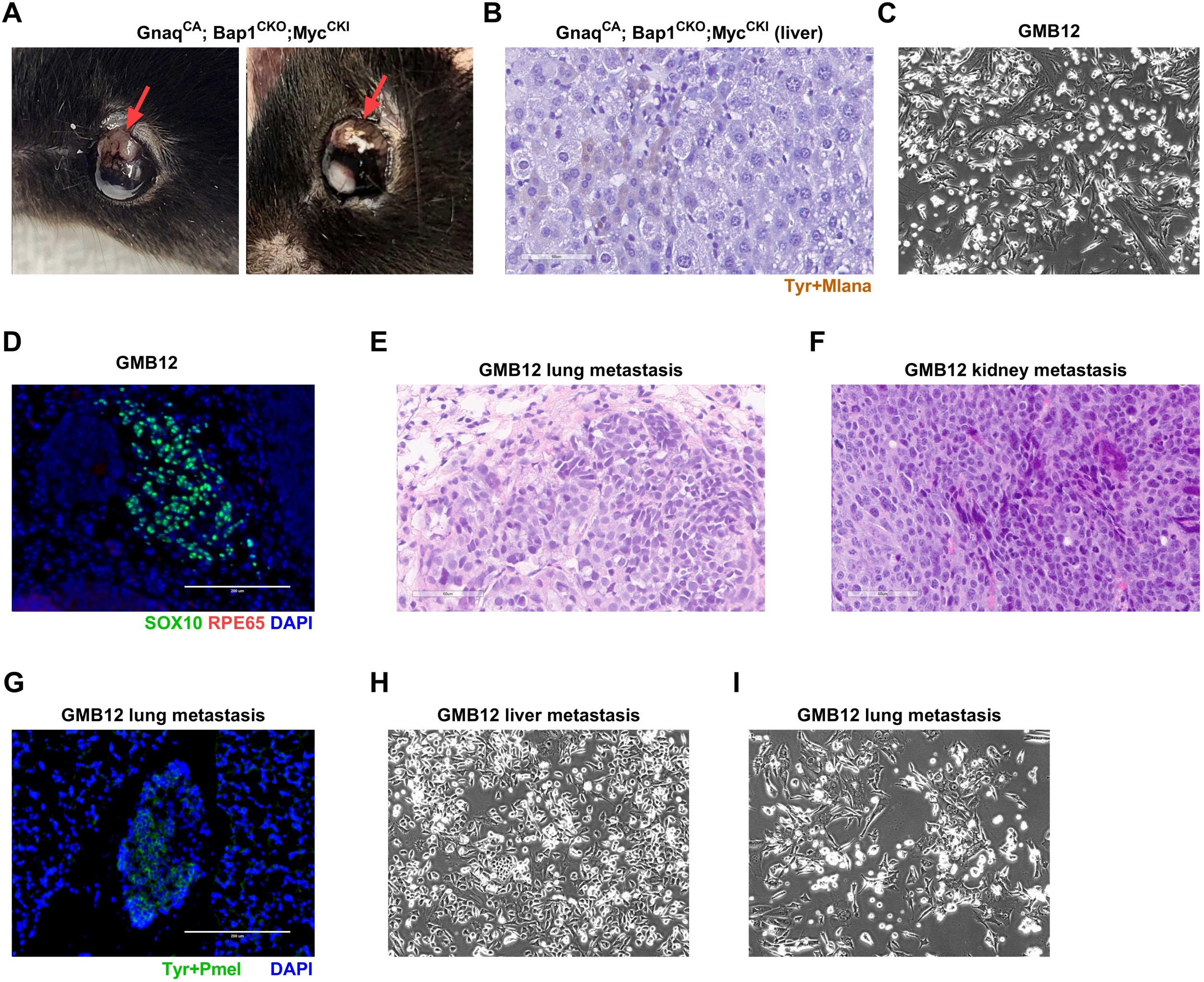
Phenotype of Gnaq^CA^; Bap1^CKO^; Myc^CKI^ mice. **A,** Representative images of eyes of Gnaq^CA^; Bap1^CKO^; Myc^CKI^ mice showing ocular proptosis and extraocular extension due to uveal melanoma development. **B,** IHC using a melanoma marker cocktail (Tyrosinase + Melan-A) in the liver of Gnaq^CA^; Bap1^CKO^; Myc^CKI^ mice. **C,** Representative image of primary UM cells GMB12 isolated from a Gnaq^CA^; Bap1^CKO^; Myc^CKI^ tumor. **D,** IHC showing SOX10 and RPE65 expression in orthotopic transplant of GMB12 cells. **E** and **F,** Representative images of H&E staining of metastasis of GMB12 cells in lung (**E**) and kidney (**F**). **G,** IHC using a melanoma marker cocktail (Tyrosinase + Melan-A) in the lung metastasis of GMB12 cells. **H** and **I,** Representative images of primary UM cells isolated from liver (**H**) and lung (**I**) metastasis of GMB12 cells.

**Figure S3.**
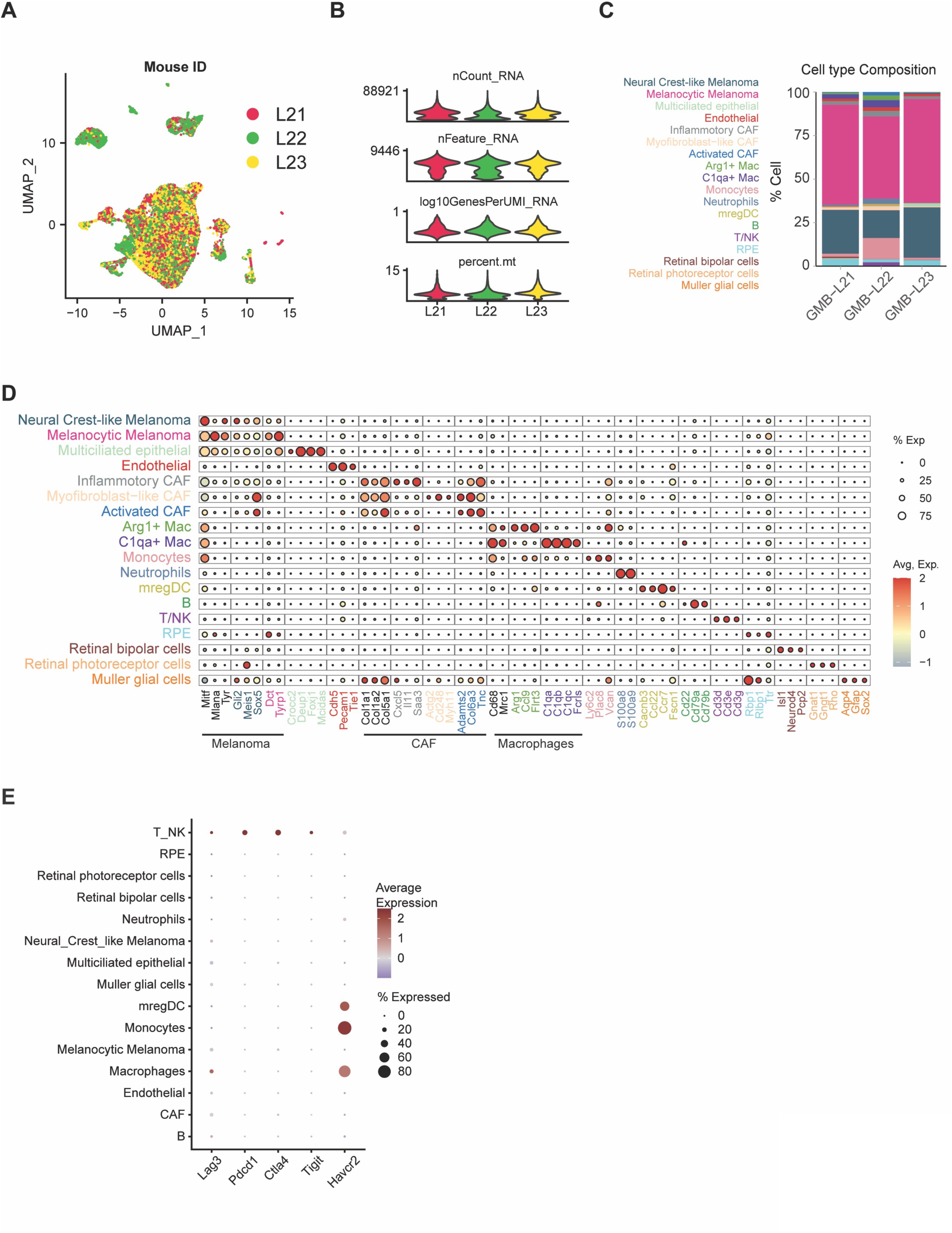
scRNA-seq characterization of murine UM composition. **A,** UMAP plot of all cell types in UM tumors from three Gnaq^CA^; Bap1^CKO^; Myc^CKI^ mice (L21, L22, L23). Cells from each mouse are labeled with different colors. **B,** Violin plots showing the distribution of key quality control metrics across samples, including total number of RNA molecules (UMI) detected per cell (nCount_RNA), number of unique genes detected per cell (nFeature_RNA), ratio of detected genes per UMI on a log10 scale (log10GenesPerUMI_RNA), percentage of transcripts mapping to mitochondrial genes (percent.mt). **C,** Proportional composition of cell types within each UM sample. **D,** Dot plot showing the expression of cell type markers, with dot color indicating the average expression levels and dot size indicating the percentage of cells in each cluster expressing the respective marker. **E,** Dot plot showing the expression of immune checkpoint genes in each cell type, with dot color indicating the average expression levels and dot size indicating the percentage of cells in each cluster expressing the respective marker.

**Figure S4.**
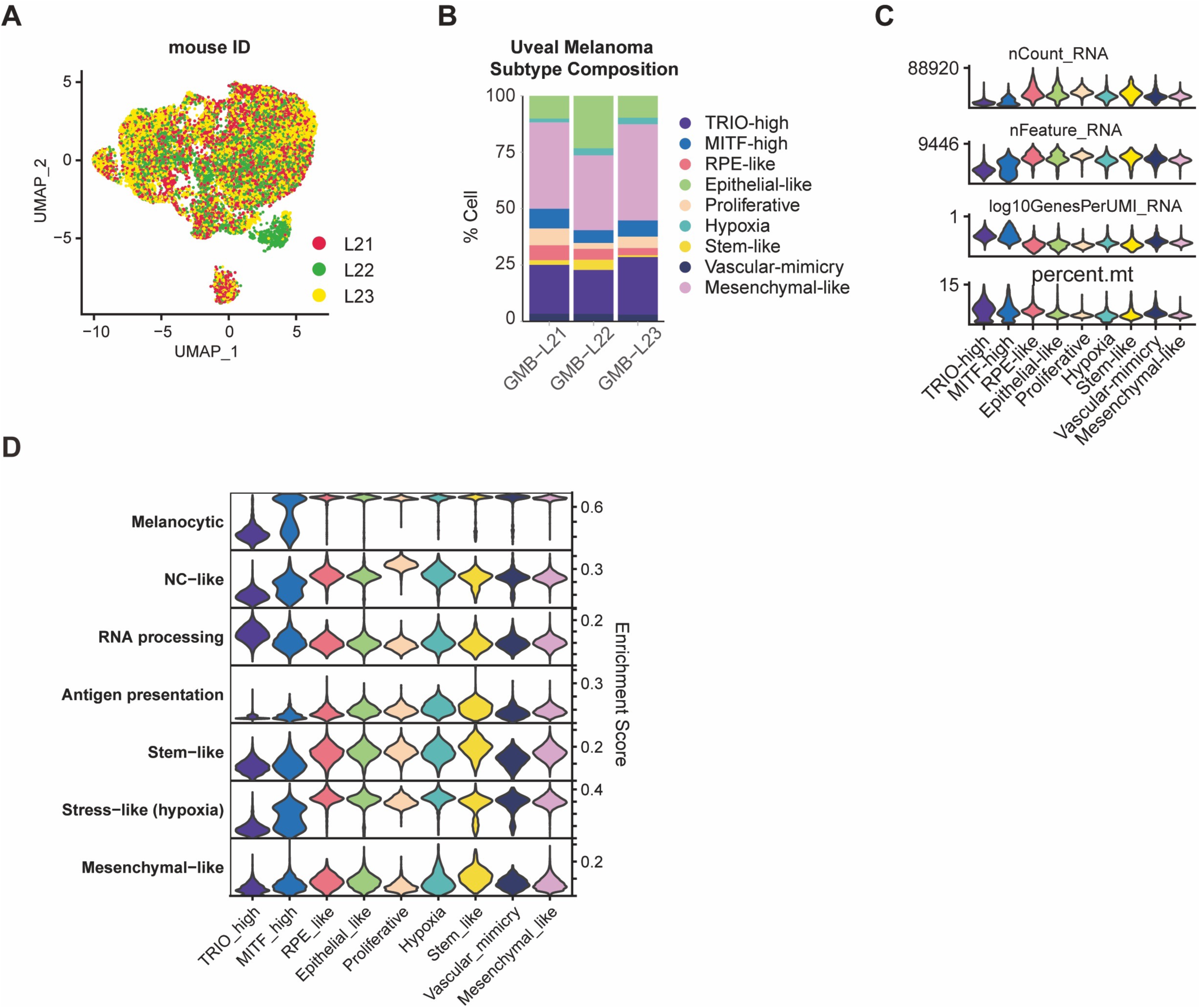
scRNA-seq characterization of phenotypic subtypes of malignant cells. **A,** UMAP plot of malignant UM cells from three Gnaq^CA^; Bap1^CKO^; Myc^CKI^ mice (L21, L22, L23). Cells from each mouse are labeled with different colors. **B,** Proportional composition of phenotypic clusters within each sample. **C,** Violin plots showing the distribution of key quality control metrics across phenotypic clusters, including total number of RNA molecules (UMI) detected per cell (nCount_RNA), number of unique genes detected per cell (nFeature_RNA), ratio of detected genes per UMI on a log10 scale (log10GenesPerUMI_RNA), percentage of transcripts mapping to mitochondrial genes (percent.mt). **D,** Violin plots showing the expression of signature defining phenotypic states of mouse cutaneous melanoma across phenotypic clusters of mouse UM.

**Figure S5.**
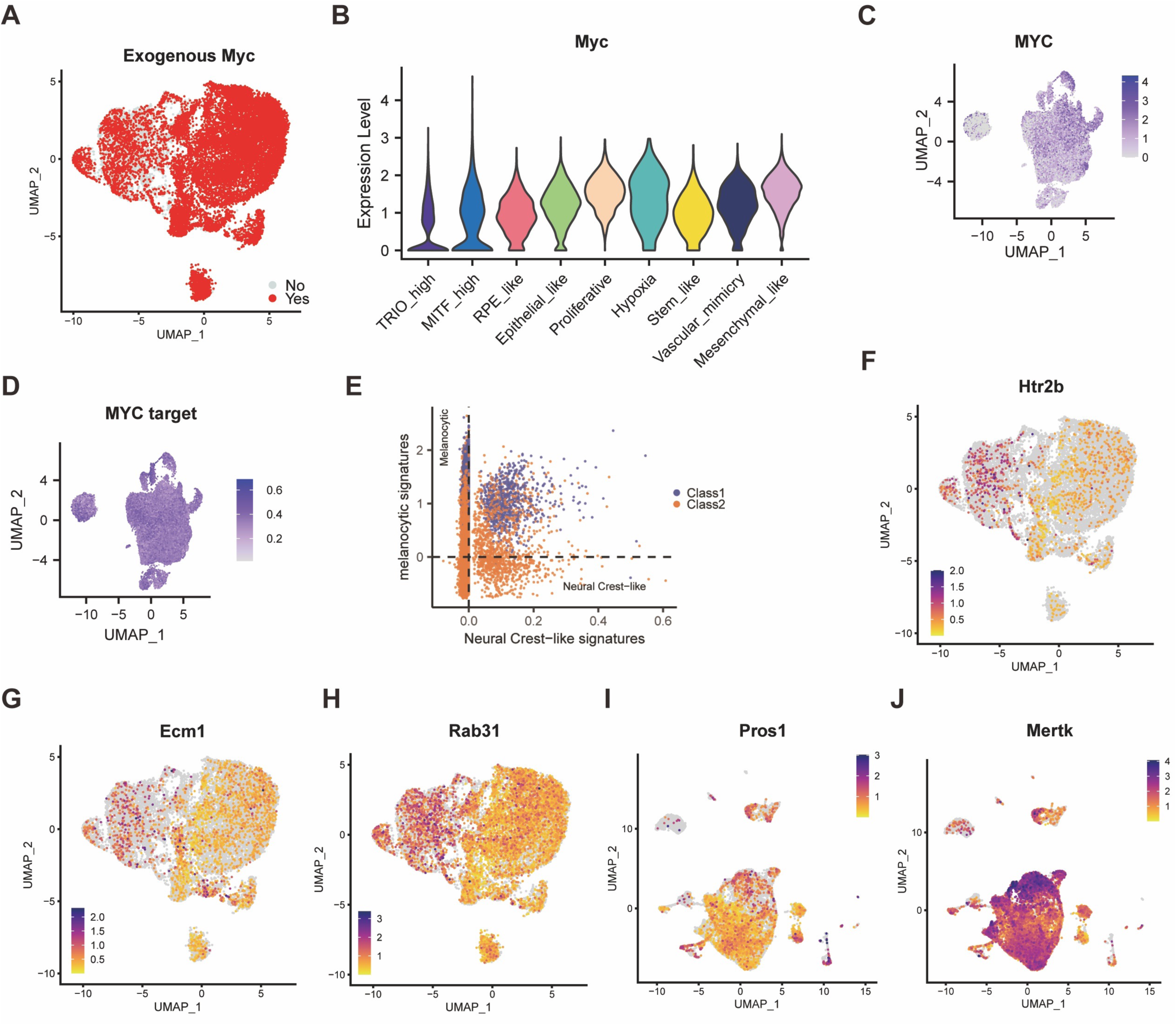
Expression of MYC and markers of human Class 2 UM. **A,** UMAP plot of UM cells from Gnaq^CA^; Bap1^CKO^; Myc^CKI^ tumors showing positivity (Yes) or negativity (No) of exogenous Myc expression. **B,** Violin plots showing the expression of endogenous Myc across phenotypic clusters of mouse UM. **C** and **D,** UMAP plots showing expression of *MYC* (**C**) and *MYC* targets (**D**) in human UM cells. **E,** Scatterplot showing positivity for the Melanocytic or Neural Crest-like signatures of Class 1 and Class 2 human UMs cells. **F-H,** UMAP plot showing expression of *Htr2b* (**F**), *Ecm1* (**G**), and *Rab31* (**H**) in malignant UM cells from Gnaq^CA^; Bap1^CKO^; Myc^CKI^ tumors. **I** and **J,** UMAP plot showing expression of *Pros1* (**I**) and *Mertk* (**J**) in all cell types from Gnaq^CA^; Bap1^CKO^; Myc^CKI^ tumors.

**Figure S6.**
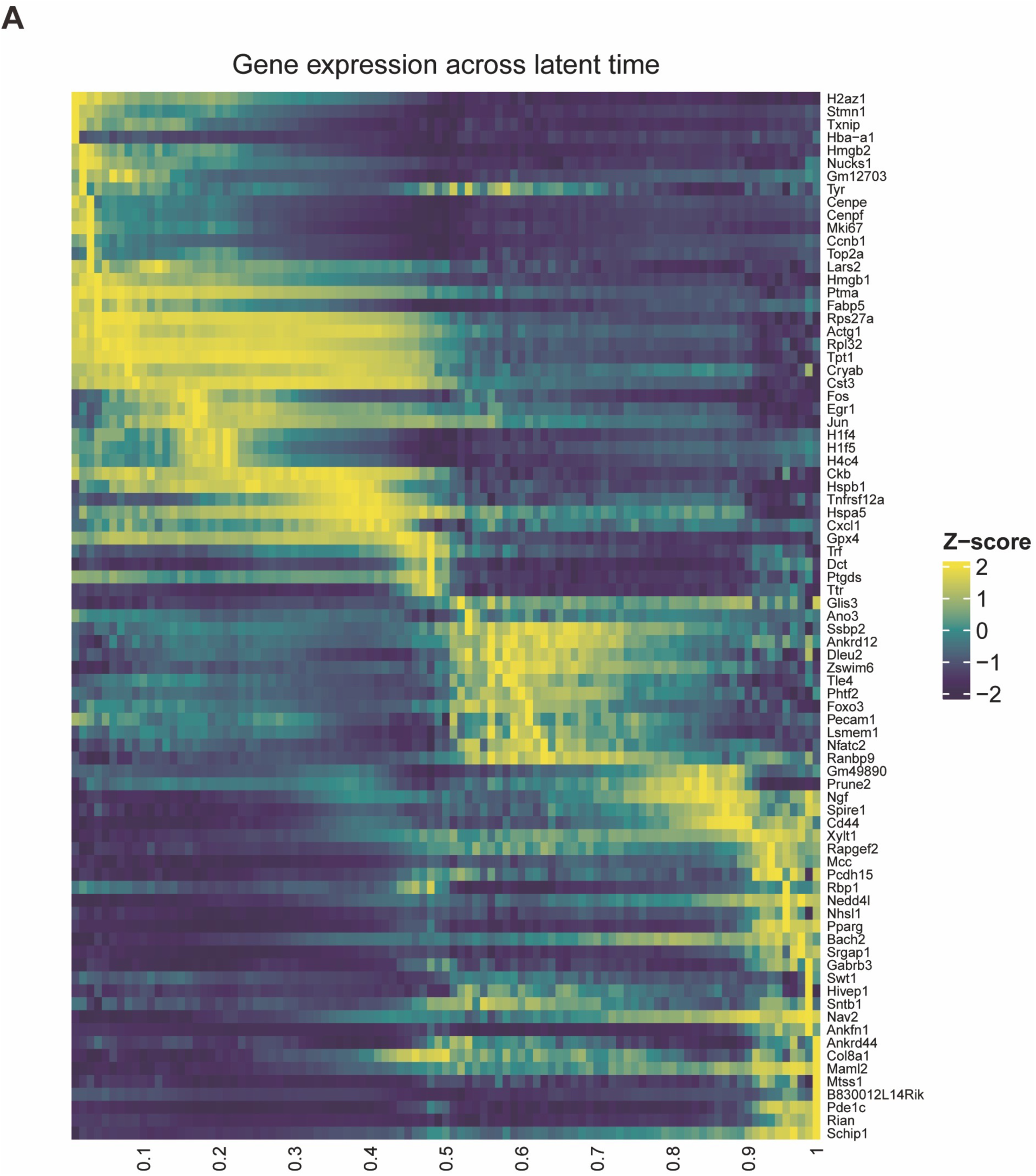
Gene expression in latent time. **A,** Expression dynamics of 81 representative genes plotted along latent pseudotime.

## AUTHOR CONTRIBUTION

X.X., J.D.L, K.S.M.S., J.W.H., and F.A.K. conceived the project and X.X., X.Y., and F.A.K. designed experiments. X.X., N.J., B.P., S.N.S.R., and V. J performed experiments. X.X., X.L., J.J.D., X.L. performed scRNA-seq analysis and X.Y. supervised bioinformatics analyses. J.K., R.L.B., J.D.L, K.S.M.S., and J.W.H provided critical insights and reagents. J.S. performed histopathological evaluations. X.L. and X.Y. performed statistical analyses. F.A.K. supervised the studies.

## ACKNOWLEDGEMENTS

We thank members of the Licht, Smalley, Harbour, and Karreth labs for helpful discussions X.X. was supported by a Melanoma Research Foundation Career Development Award (1068914). J.D.L, K.S.M.S., and J.W.H. were supported in part by R01CA256193 from the NCI. J.W.H. was also supported a Cancer Prevention and Research Institute of Texas Recruitment of Established Investigator Award (RR220010), NCI Cancer Center Support Grant (P30CA142543) to University of Texas Southwestern Simmons Comprehensive Cancer Center, NEU Core Grant (P30EY030413) to University of Texas Southwestern Department of Ophthalmology, and Research to Prevent Blindness, Inc. Challenge Grant to University of Texas Southwestern Department of Ophthalmology. F.A.K. was supported by grants from the Melanoma Research Alliance (https://doi.org/10.48050/pc.gr.154417) and the Department of Defense Melanoma Research Program (ME230182). This work was also supported by the Gene Targeting Core, Bioinformatics and Biostatistics Shared Resource, Molecular Genomics Core, and Analytical Microscopy Core, which are funded in part by Moffitt’s Cancer Center Support Grant (P30CA076292). K.S.M.S. receives funding from Revolution Medicines and Neogene and J.W.H. receives royalties from Washington University for IP that was licensed to Castle Biosciences related to prognostic testing in uveal melanoma, unrelated to this work. J.W.H. is a consultant for Castle Biosciences.

